# CAPO: Compute-Aware Automated Protein-Model Optimization

**DOI:** 10.64898/2026.10.12.758406

**Authors:** Théo Schifferli, Julián García Pardiñas, Edward B. Irvine, Mary-Anne Hartley, Thomas Bikias

## Abstract

Protein models such as Evolutionary Scale Modeling (ESM), AlphaFold and Boltz have transformed computational protein science, yet a pretrained model rarely transfers to a particular laboratory’s assay without adaptation to its data and experimental setting, and that adaptation is technically demanding and spends real compute budget. Existing agentic systems can automate model training, yet typically treat fine-tuning as a reliable tool call: feasibility and cost are estimated rather than measured, and infeasibility is discovered only after compute has been spent. We formulate protein-model adaptation as a gated decision problem: given a task, raw data and a compute budget *c*_max_, a system should decide *whether and how* to adapt a model before committing expensive computation. We introduce CAPO (Compute-Aware Automated Protein-Model Optimization), a multi-agent system built around one rule: gate before you train. CAPO measures feasibility on the target accelerator, projects cost against the available budget, runs a canary inside the actual training loop and maintains an auditable artifact trail; critically, it can decline an unaffordable adaptation and select a cheaper alternative. We evaluate CAPO, without system changes, on three protein-modeling tasks: (i) SARS-CoV-2 host-range prediction, (ii) hit-to-lead affinity ranking and (iii) temporal variant-effect reclassification. On host range, CAPO partially fine-tuned an 8-million-parameter ESM-2 model to a species-averaged MCC of 0.826, compared with 0.743 for a hand-designed expert transformer; it matched a general coding agent while using 57.4% less GPU time and reduced failed training trials from four to zero. For hit-to-lead affinity ranking, CAPO rejected Boltz-2 fine-tuning and instead trained an assay-calibrated lightweight head over frozen representations, improving ranking on 11 of 15 held-out targets in 1.68 GPU-hours. On temporally held-out ClinVar variants, rank-4 low-rank adaptation of ESM-2 reached an MCC of 0.478, compared with 0.252 for a one-hot baseline and 0.000 for the frozen zero-shot score. These results come from single runs, without a gate ablation and with one generalist-agent comparison. Within those limits, they show that measured admission control can make protein-model adaptation not only automated but selective: deciding what to compute, what not to compute and why.

## 1 Introduction

Protein foundation models have changed what a single laboratory can compute: sequence models such as ESM-2, and structure or co-folding models such as AlphaFold and Boltz, turn amino-acid sequences and related molecular inputs into predictions that would otherwise require extensive experimental work [1–3]. What they do not do is answer a specific laboratory’s question: practical protein engineering needs predictions aligned to a particular assay and setting, and reaching them almost always requires adapting a pretrained representation to task-specific measurements [4–7]. Pretraining is paid once by a model’s authors, and inference on a released checkpoint is now cheap. The dominant cost of applying these models is neither: it is the fragile, per-assay adaptation between a raw experiment and a model a scientist can trust. Raw assay output must become a valid sequence– label dataset; a backbone must be chosen, and with it an adaptation technique, whether full or partial fine-tuning, low-rank adaptation or a head over frozen representations, whose right setting depends on the dataset rather than the model [7]; a pipeline must be written and run on a paid accelerator; and the result must be evaluated so that near-duplicate sequences shared between train and test do not flatter it. Each stage fails in ways that are hard to see, and doing it well demands both biological and machine-learning expertise.

### Why current agent harnesses are not enough

Tool-using agents already plan, generate code, launch jobs and report costs [8–11], and automated machine-learning agents build competitive models on biomedical benchmarks [12–14]. Two of their assumptions become load-bearing once the tool is a fine-tuning run. First, that tools are cheap and reliable. A fine-tuning run is neither: it consumes hours of paid accelerator time and can exhaust memory, diverge or silently underfit, so the decision to spend is made from estimates or discovered by failing, and a bad plan is paid for in full before it is detected. Second, that successful execution implies scientific correctness. An agent that finishes and reports metrics may still have leaked its test set or corrupted its inputs, and its own confidence is a poor guide to whether the model survives held-out data [14]. Recent analyses of agentic drug-discovery systems name the gap directly: protein foundation models are still not supported as trainable, first-class components [15].

### Adaptation as a gated decision problem

We therefore reframe the task. Given a biological objective in natural language, raw experimental data and a compute budget *c*_max_, an agentic system should determine whether a pretrained model is to be adapted at all, and by which technique, before committing expensive computation. This makes admission control, rather than code generation, the organising principle of the harness: every expensive action is admitted only against a measurement taken on the hardware that will run it, and the system must be able to decline. We call the rule “gate before you train”, and implement it in CAPO (Compute-Aware Automated Protein-Model Optimization).

### Contributions

- **A harness whose organising principle is measured admission control.** We formalise a feasibility probe on the target accelerator, a cost projection, a tolerance-band admission gate that may reject, a canary re-measurement inside the real training loop, and reconciliation of projection against realised cost (Section 3). To our knowledge no prior system for protein-model adaptation combines measured feasibility, explicit budget admission, leakage-aware evaluation and the ability to decline.
- **A controlled generalist-versus-specialist comparison.** On identical raw paired-end sequencing data, task text and USD 50 budget, a domain-specific harness and a general coding agent are compared on accuracy and on process cost: accuracy is matched, GPU time falls by 57.4%, cost per successful run by 32.5% and failed trials from four to zero, and upstream of any model the general agent’s raw-read curation scores 0.00% against a blinded golden set while appearing to succeed (Section 5.1). This is one tightly controlled task, not three head-to-heads, and its halves disagree: on modelling the agent ties; on the data engineering before it, it fails completely and quietly.
- **Evidence that one unchanged system selects a different adaptation technique per task, including refusing to spend.** The same code partially fine-tunes ESM-2 on nine million labelled reads, declines to fine-tune Boltz-2 and fits an assay-calibrated head over the frozen backbone instead, and applies rank-4 low-rank adaptation on a small, temporally split clinical benchmark, with null results surfaced rather than dropped (Sections 5.2 and 5.3).

## 2 Related work

Tool-using agents established the control structure CAPO’s orchestrator inherits, from interleaved reasoning and action [8] to model registries as callable tools [9] and composed domain tools in scientific workflows [10]; automated machine-learning agents extend it to model building, including on biomedical data with stricter validation [12–14]; and general-purpose engineering agents now research, write and ship machine-learning code end to end against a public model and dataset ecosystem [11, 16]. All decide what to call rather than whether calling it is affordable and feasible on the hardware at hand, so infeasibility is discovered by failing.

In protein engineering, AutoProteinEngine is the closest prior work, bringing natural-language-driven automated machine learning with model selection and hyperparameter search [17], while platforms such as Tamarind Bio and Biomni Lab make a broad library of protein tools, and a fine-tuning menu over it, usable without machine-learning training [18, 19]. We read these as complementary rather than competing: they widen access to the tools, whereas CAPO is concerned with which tool to spend on, and whether to spend at all. Both begin from a prepared dataset rather than a raw assay, and both present adaptation as a menu choice, with the reasoning behind a configuration and its price left implicit. CAPO does not hand the user a menu. It reads the raw assay, recommends one strategy and the measured evidence for it, prices that strategy on the accelerator that will run it, and writes an artifact for every decision, so what the user receives is an auditable, costed recommendation that can be checked rather than trusted.

The protein engineering literature says what such a harness must encode: performance under a fixed budget depends strongly on how model size and data are allocated [20]; parameter-efficient methods recover much of the benefit of full fine-tuning at a fraction of the cost [6]; whether feature extraction, low-rank adaptation or full fine-tuning is right is dataset-dependent [7]; and agentic protein workflows need domain tools and quantitative validation to produce grounded outputs [21]. Technique selection is therefore knowledge a harness must encode, not a default it may assume. CAPO’s delta is the combination of feasibility measured on the target accelerator rather than estimated, a budget admission decision that may decline, leakage-aware evaluation inside the pipeline rather than after it, and reconciliation of projection against realised cost; to our knowledge no prior system for protein-model adaptation has all four.

## 3 CAPO: an auditable, reproducible, compute-aware system

**Figure 1:**
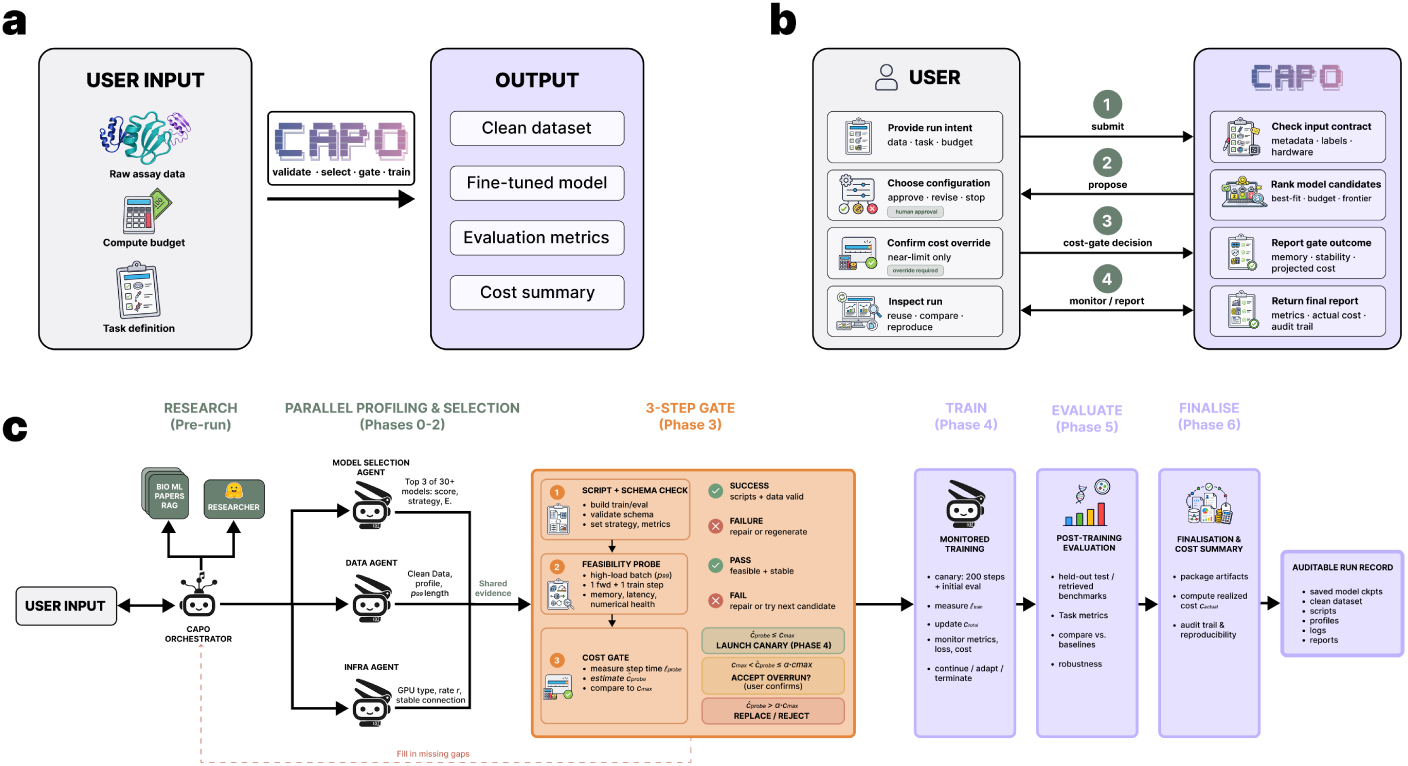
CAPO. (a) Raw assay, task definition and budget *c*_max_ in; processed dataset, adapted model, evaluation and cost reconciliation out. (b) The user holds two decisions: confirming the backbone, and accepting or rejecting a projected overrun. (c) Pre-run and Phase 0–2 agents profile the data, provision hardware and rank backbones concurrently; their evidence feeds the **three-step gate of Phase 3**, where a script and schema check, a feasibility probe on the target accelerator and a cost gate against *c*_max_ must all pass before any GPU-hour is committed.

CAPO is a multi-agent system built on the Claude Agent SDK. One orchestrator, on Opus 4.8, drives each run and delegates narrowly scoped work to eleven specialised agents, each pinned to a fixed model, a written system prompt, a restricted toolset and an execution limit, communicating only through persistent artifacts in a shared run directory (Appendix C). Compute-awareness operates at two levels: feasibility and cost checks stop a configuration reaching a long run unless it fits the hardware and the budget, and model capacity is matched to the risk of each role, so a read-only health check running every minute uses Haiku 4.5 while open-ended planning, code synthesis and failure recovery use the largest model. Auditability is enforced, not encouraged: every decision is written to disk as a structured artifact, and a missing or unparseable artifact is a hard failure. Reproducibility cannot mean an identical reasoning path on every run; it means a run preserves its generated code, configurations, launch commands and model identifiers, so the same inputs reproduce the same decision path.

### 3.1 Before the gate: evidence, not defaults

Two advisory agents open a run, both overridden by current-run measurements on conflict so that a stale prior cannot quietly corrupt a fresh run: a **memory consultant** giving CAPO episodic memory over a cross-run index of past run reports, and a **Hugging Face researcher** proposing datasets, the field-standard metric and an initial hyperparameter regime, verifying every identifier against the model hub in-session and reporting anything unconfirmed as unknown. Three characterisation agents then run concurrently. The **infrastructure agent** provisions the accelerator and fixes the hourly rate *r*, shared unchanged with the Phase 3 gate and the Phase 6 reconciliation so projected and realised cost are computed under one resource model. The **data profiler** decides whether the input can support a valid sequence–label dataset at all, classifying it model-ready, transformable or incompatible; a transformable input is accepted only with assay metadata specifying the extraction procedure, target region, reading frame and label rule, and one lacking it is rejected before model selection, because launching on ill-defined data is the most expensive mistake the system can prevent. Its profile records the task formulation, label statistics, preprocessing decisions and sequence-length percentiles; the *p*_99_ length defines the high-load batch the Phase 3 probe must survive. The **model selector** ranks a curated registry of 30 protein models, from ESM-2 at 8M to 15B parameters through to structure and affinity predictors such as ESMFold and Boltz-2 [1, 22, 23], by a transparent weighted score *s* = 0.40 *f_t_* + 0.20 *f_m_* + 0.15 *f_c_* + 0.10 *f_o_* + 0.10 *f_a_* + 0.05 *f_d_*over six factors in [0, 1] scoring task fit, modality fit, compute fit, openness, maturity and deployment readiness, with penalties for gated access, over-budget compute or a non-commercial licence. It returns three candidates spanning the cost–capability trade-off and the user confirms one, but this score only prioritises candidates for probing: admission depends on the measured gates below. The adaptation technique is initialised from a deliberately conservative prior keyed on label count (Appendix B), then revised against task, data, model and budget evidence: a starting point, not a rule.

### 3.2 The gate: probe, cost admission, canary

#### Feasibility probe

The orchestrator first synthesises task-specific training and evaluation code rather than filling a generic template, which lets the same system handle classification, regression and per-residue tasks with the correct loss, head and metrics; a shared train-step factory is imported by both the probe and the training script, so the mixed-precision backward pass the probe exercises is the one full training will run. The probe answers one narrow question before any reasoning about cost: can this configuration run on this accelerator? It builds a demanding batch from sequences near the profiled *p*_99_ length and runs inference and one full optimisation step while measuring peak memory, step latency and numerical stability. Failures are diagnosed by cause and handled by a bounded repair ladder (batch size, precision, dataset parsing, code regeneration, escalation to a repair critic); if no candidate passes, the run terminates with a diagnostic report. A memory or numerical failure found five minutes into a probe is free; five hours into training it is not. The probe also yields the first honest cost estimate: with *E* the resolved epoch count, *N* the training-set size, *B*_eff_ the effective batch size after gradient accumulation and *ℓ*_probe_ the measured duration of one complete optimisation step,

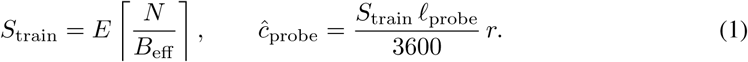

#### Cost gate

The gate turns *ĉ*_probe_ into an admission decision, with *α* = 1 + *τ* for a user-defined tolerance *τ ∈* [0 1]:

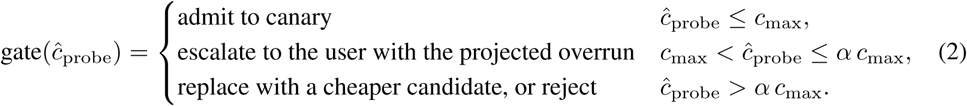

Only configurations simultaneously executable, numerically stable and affordable are admitted, and the human is kept in the loop at exactly the boundary case where a small overrun might be worth accepting. The third branch is what makes this a decision procedure rather than a budget flag: the system may conclude that the requested adaptation should not happen, and propose a cheaper one.

#### Monitored canary

An admitted candidate opens with a canary of *T*_canary_ = 200 optimisation steps inside the real training loop. The probe establishes that a backward pass can execute; the canary tests whether the model actually learns, aborting on a non-finite loss, an exploding gradient norm or a loss slope that fails to decrease. It also replaces probe timing with a measurement from the launched script, capturing data loading, logging, scheduler and optimiser behaviour a probe cannot. With *t*_canary_ the canary wall-clock time, *q*_eval_ the evaluation interval in steps and *t*_eval_ the duration of the first evaluation pass,

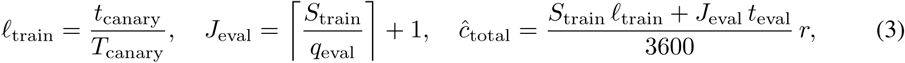

where the canary steps are already counted in *S*_train_. Training continues only if the canary is healthy and *ĉ*_total_ is still within budget; otherwise a bounded correction is applied, and when that is insufficient a recovery agent diagnoses the failure and its proposed fix must pass a new canary before training resumes.

### 3.3 Monitoring, finalisation and reconciliation

A concurrent **health monitor** watches the live run read-only, one remote round-trip per check, so it is cheap to run often: it can recommend termination on repeated divergence, non-finite values, exploding gradients, stagnation or budget drift, while tolerating a transient loss spike that leaves validation metrics intact. Evaluation is on data excluded from both optimisation and online model selection, with homologous sequences clustered and assigned at cluster level when no held-out split is provided (Appendix B). The **finaliser** then writes an evidence-backed report in which each claim derives from a specific artifact and any unavailable value is recorded as unknown rather than fabricated; runs that terminate early are finalised in the same format, so a failed run is as inspectable as a successful one. Realised cost comes from measured wall-clock accelerator runtime *t*_wall_ at the rate *r* the gate used, *c*_actual_ = (*t*_wall_*/*3600) *r*, and is reported beside *ĉ*_probe_ and *ĉ*_total_, so a user inspects not only the final cost but the accuracy of the system’s own forecasts; across the three studies the gate overestimated realised GPU cost by 2.1*×* to 3.2*×* (Appendix G), an error in the direction that keeps a run inside its budget. The report frontmatter is appended to the cross-run index, becoming a prior for the next run’s memory consultant.

## 4 Evaluation design

Three real biological tasks stress different parts of the harness (Table 1). Every run received the same USD 50 budget and was executed end to end by the same unmodified system from a single natural-language request, reproduced verbatim in Appendix J; construction, splits and baselines are in Appendices D to F.

**Table 1:** The three evaluation tasks, stressing respectively data engineering at scale, admission control against an unaffordable adaptation, and leakage-aware evaluation on a small, skewed clinical dataset. Budget: USD 50 each.

| Task | Data | Primary metric | Baselines |
| --- | --- | --- | --- |
| Host range | Raw paired-end reads; $\sim 9.5\text{M}$ labelled examples at 201 aa | Species-averaged MCC | General coding agent; expert transformer; frozen-embedding MLP and probe |
| Hit-to-lead | FEP+ benchmark; 644 co-folds, 15 scorable series | Spearman, log-Pearson, RMSE | Raw and calibrated Boltz-2; pooled kernel-ridge regressor |
| Variant effect | ClinVar 2020 to present; eight-gene panel; 420 reclassified VUS | MCC ($n=420$) | One-hot logistic regression; frozen ESM-2 zero-shot |

Host range is the only task starting from genuinely raw instrument output: turning a yeast-surface-display screen of an Omicron BA.1 RBD mutagenesis library into supervised examples means merging read pairs, trimming adapters and flanks, filtering low-quality and rare reads, and translating each survivor into its canonical 201-residue domain [24, 25]; keeping only single-label variants leaves supervision fragmented and strongly imbalanced (3.2% binders for rat, 51.4% for mink). That curation step is scored blind, on two host libraries not used for modelling, against a golden set withheld from both systems; the human comparator is a per-residue transformer refined over several iterations for this assay (Appendix D). Hit-to-lead uses Boltz-2, which is far too large to fine-tune inside a USD 50 budget, and that is the point; each FEP+ series [26, 27] is split by a temporal proxy on compound numbering with a Murcko-scaffold fallback so no scaffold straddles the two batches, held-out labels are touched exactly once at final scoring, and Boltz-2’s weights are never updated by any of the four compared methods. Variant effect forbids lookahead by construction: models are fitted on variants already definitive in the January 2020 ClinVar release and tested only on variants uncertain in 2020 and since reclassified [28], all three receiving identical, strictly sequence-only inputs, namely a 401-residue reference window and its mutated counterpart, with no gene identity, residue number, phenotype or external pathogenicity score. Eight proteins are too few for a homology-clustered split, so leakage control rests on the temporal split, a position-grouped internal split and an explicit position-novel and region-novel audit, with all thresholds locked on validation data before the temporal cohort is opened.

## 5 Results

Results follow the task order of Table 1 and are interpreted in Section 6; Table 4 in the appendix aggregates the three runs.

### 5.1 Host range: generalist versus specialist, measured twice

#### Upstream, the general agent completes and scores zero

Scored blind against an independently curated golden set, CAPO recovered canonical full-length receptor-binding domains at 97.6% agreement on the pangolin library of 14.4M sequences (USD 9.91) and 99.83% on the possum library of 2.61M sequences (USD 6.88). Given the identical raw reads, the general coding agent stopped at read merging and returned read-window fragments of 79 to 167 residues that never span the domain, so its agreement was 0.00% on both, at USD 25.96 and USD 7.09 (Table 8). Agreement requires exact identity with a full-length domain, which a fragment cannot satisfy (Appendix D). Neither agent run crashed or declined the task: both terminated normally and returned plausibly formatted sequence files.

#### Downstream, accuracy is matched and process cost is not

With backbone and adaptation regime fixed in advance, CAPO adapted ESM-2 8M [1] by unfreezing its last four transformer layers under a multi-label head, training five epochs in bfloat16 with AdamW. It reached a species-averaged MCC of 0.826, against 0.743 for the expert’s hand-designed, iteratively tuned transformer, a gain of 0.083 or 11.2% relative, and 0.812 for the general coding agent, for which no confidence intervals are available (Figure 2). Per species CAPO matched or exceeded the agent on 7 of 10 orthologs, with the larger gaps on the lower-MCC species. On process cost, CAPO reached its reported model with 57.4% less GPU time, 32.5% less cost per successful run, and zero failed training trials against the agent’s four.

**Figure 2:**
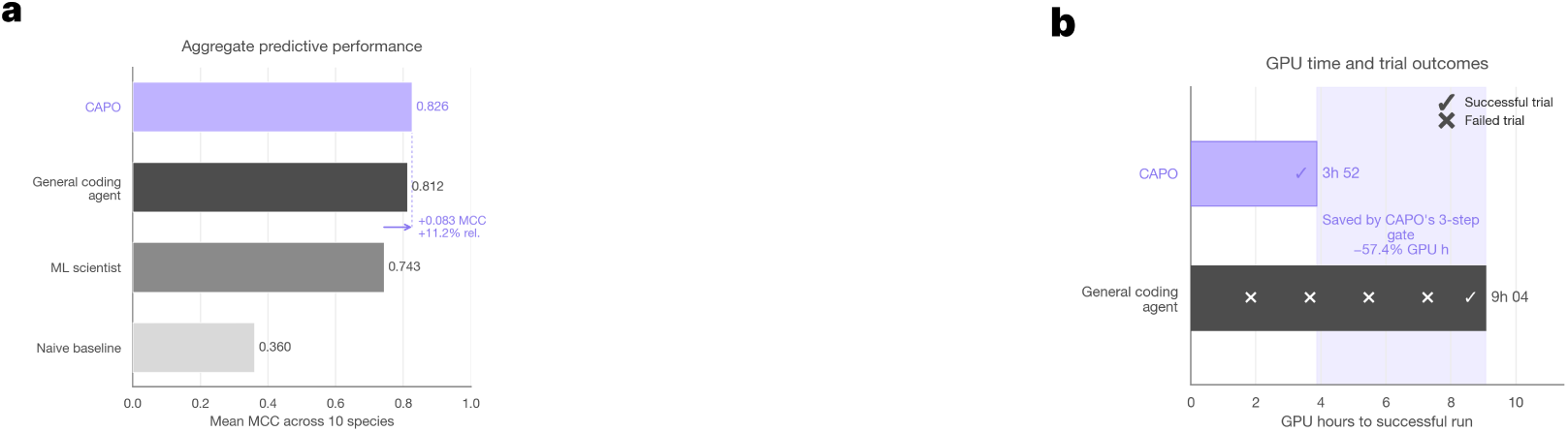
Accuracy ties; process cost does not. Left: species-averaged MCC; CAPO exceeds the hand-designed expert transformer (labelled “ML scientist”) by +0.083 and matches the general coding agent. Right: GPU hours to a successful run, trials successful (✓) or failed (*×*). See Figures 4 and 5.

### 5.2 Hit-to-lead: the declined fine-tune

#### The gate declined the fine-tune

Asked to improve hit-to-lead ranking with Boltz-2 under the same USD 50 budget, CAPO did not fine-tune it. The projected cost of a weight update to Boltz-2 fell in the third branch of Equation (2), so the configuration was replaced rather than escalated to the user. The system kept Boltz-2 frozen, reused 644 cached co-folds and fitted a per-target assay-calibrated affinity head on its outputs [29].

#### The cheaper strategy leads every comparator

Across the 15 FEP+ series with a scorable held-out batch the head led on every reported metric (Table 2): Spearman 0.700 against 0.594 for raw Boltz-2, log-Pearson 0.740 against 0.623, and RMSE 0.559 log_10_ *K_d_* units against 1.061 for raw Boltz-2 and 0.610 for its calibrated form, clearing a pre-registered success threshold of 0.35. Per target it improved log-Pearson on 11 of 15 series over raw Boltz-2 (median +0.086, paired Wilcoxon *p* = 0.073) and on 12 of 15 over the pooled classical regressor (median +0.083, *p* = 0.015); the two baselines did not separate from each other (median difference 0.005, *p* = 0.68). The whole run consumed 1.68 GPU-hours at USD 1.99 per hour, a compute cost of USD 3.35. An ablation attributes the ranking gain to the leave-one-out-gated QSAR blend and almost all of the error reduction to calibration (Appendix E).

**Table 2:** Ranking quality on the 15 scored FEP+ series, averaged over targets; Boltz-2’s weights are frozen in all four methods. Raw Boltz-2’s error is on a scale mismatched to the potency labels, so the fair error comparison is against calibrated Boltz-2; best value per column in bold.

| Method | Spearman ↑ | log-Pearson ↑ | RMSE ($\log_{10} K_d$) ↓ |
| --- | --- | --- | --- |
| Raw Boltz-2 | 0.594 | 0.623 | 1.061 |
| Calibrated Boltz-2 | 0.594 | 0.623 | 0.610 |
| Pooled classical ML | 0.616 | 0.642 | 0.667 |
| **CAPO head (frozen backbone)** | **0.700** | **0.740** | **0.559** |

**Figure 3:**
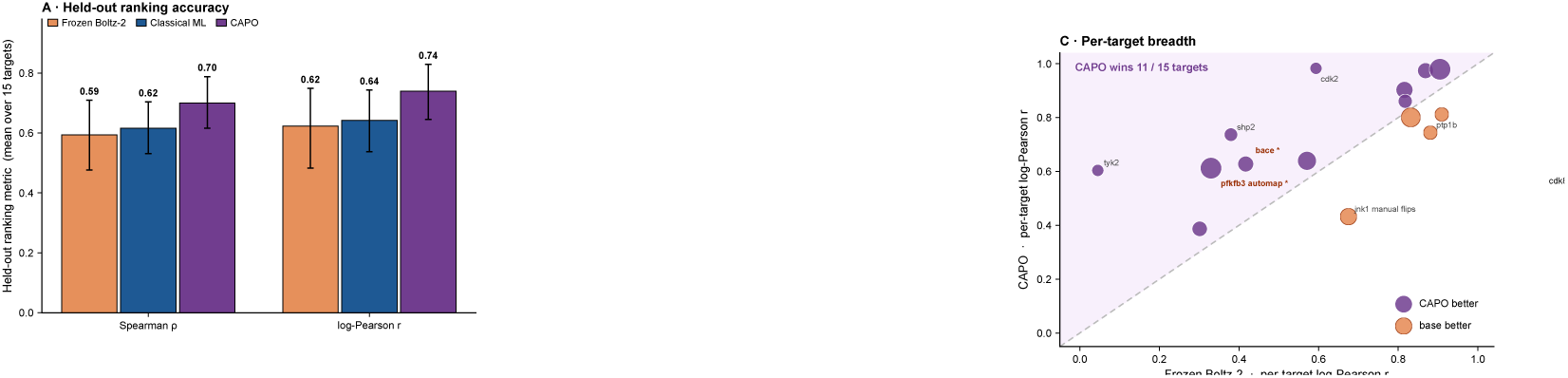
An assay-calibrated head over a frozen co-folding model, chosen because fine-tuning was refused. Left: held-out ranking accuracy over the 15 scored series, 95% bootstrap intervals. Right: per-target log-Pearson; points above the diagonal are targets CAPO ranks better than frozen Boltz-2, stars the two where Boltz-2 ranks worst.

#### Coverage and null results

The pre-registered hardest-case contrast, targets simultaneously hard for the physics model and for a classical regressor, was empty on this benchmark, so the reported average is over the 15 scorable series. On the two targets where Boltz-2’s own affinity ranks worst, BACE and PFKFB3, mean raw log-Pearson rose from 0.37 to 0.62, with the classical baseline also competitive there. The FEP+ physics anchor mapped none of the 214 held-out ligands, and 14 of the 29 co-folded targets carried held-out batches too small or too single-scaffold to rank.

### 5.3 Variant effect: leakage-aware temporal adaptation

#### Thresholds locked before the temporal cohort was opened

CAPO ran this study end to end on one remote A100 (40 GB SXM4) without human intervention: it profiled the raw ClinVar tables, built the three locked cohorts, passed the probe and cost gate at a projected USD 4.26, fine-tuned three seeds under rank-4 low-rank adaptation [30] of ESM-2 150M, locked calibration and thresholds before opening the temporal cohort, and wrote its own report. The leakage audit found zero intersection between the training and temporal cohorts. The run realised USD 29.76 in about 1.5 hours against USD 4.26 projected.

#### Adaptation improves discrimination over both baselines

On the temporal cohort of 420 variants uncertain in 2020 and since reclassified, adaptation reached an MCC of 0.478, against 0.252 for the one-hot logistic floor and 0.000 for the frozen zero-shot score (Table 3). Over 10,000 paired stratified bootstrap samples the deltas were +0.226 with a 95% interval [0.120, 0.334] over one-hot and +0.478 with [0.403, 0.551] over zero-shot, with McNemar’s test at *p* = 2.3 *×* 10*^−^*^6^ and *p* = 3.9 *×* 10*^−^*^18^. The frozen zero-shot model reached an AUROC of 0.878 while scoring an MCC of 0.000 at its fixed threshold; adaptation raised AUROC to 0.892 and balanced accuracy from 0.500 to 0.721.

**Table 3:** Temporal reclassified-VUS benchmark (Cohort B, *n*=420). All three models receive identical, strictly sequence-only paired windows; best per column in bold.

| Model | MCC ↑ | AUROC ↑ | AUPRC ↑ | Bal. acc. ↑ | ECE ↓ |
| --- | --- | --- | --- | --- | --- |
| One-hot logistic | 0.252 | 0.643 | 0.606 | 0.599 | 0.198 |
| ESM-2 zero-shot | 0.000 | 0.878 | 0.884 | 0.500 | 0.312 |
| **ESM-2 + LoRA ($r=4$)** | **0.478** | **0.892** | **0.888** | **0.721** | **0.190** |

#### Novelty audit, calibration and coverage

A novelty audit split a TP53 saturation scan by distance to the training data. On the 303 position-novel variants, with no training variant at the same residue, the advantage over the frozen model was +0.424 ([0.332, 0.512]); on the 49 region-novel variants, excluding any training variant within *±*10 residues, it was +0.213 with an interval containing zero (Appendix F). Only 446 of the 2,897 historical definitive labels available for development were benign (Table 9). The adapted model recorded the lowest expected calibration error of the three at 0.190. CAPO also scored the 1,000 variants of Cohort C that remain uncertain today, for which no ground truth exists.

## 6 Discussion

### One system, three techniques

The argument of this paper is what the three runs have in common: the same code, the same gates and the same USD 50 budget produced three different adaptation techniques, because the measured evidence differed. Abundant labels and a small backbone bought partial fine-tuning of ESM-2 8M; a large backbone under a small budget bought a refusal, and a fall back to an assay-calibrated head over frozen representations; few labels and binding leakage bought rank-4 low-rank adaptation of ESM-2 150M under a temporal split. Two of the three differ from the conservative label-count prior that initialised them (Appendix B): measurement overriding a default rather than confirming it. They were not chosen by us and not drawn from a per-task menu; they are what the gate admitted (Table 4).

### What measured admission control buys

Three things. It converts a class of expensive failures into cheap ones: each of the four trials the general agent burned failed a feasibility or affordability check that was never applied, and refusing them before launch rather than on an accelerator is where the 57.4% GPU-time and 32.5% cost reductions come from. It makes not spending an available action: at USD 1.99 per GPU-hour the whole hit-to-lead budget buys about 25 GPU-hours, nowhere near a weight update to a model of Boltz-2’s size, so the overrun was not one a user could sensibly be asked to accept, and the fallback the gate chose instead still improved ranking on 11 of 15 targets for 1.68 GPU-hours. Most automated pipelines know only how to spend well; a procedure that can return no covers the case where the budget does not buy the intervention. And it makes the system’s own forecasts auditable: projection and reconciliation share one resource model, so a mis-priced run shows up in the run record, not in a cloud bill.

### Where the gate is wrong

Admission control is only as good as its price model, and ours is wrong in two directions the run record makes visible. Against realised GPU cost it is pessimistic, over-projecting by 2.1*×* to 3.2*×* (Appendix G), because *ℓ*_probe_ in Equation (1) is measured on a *p*_99_-length batch and then held for the whole schedule; its errors are false refusals rather than budget overruns, the safe direction, but an affordable run can still be declined. Against the realised cost of the whole run it is optimistic, pricing the training loop and not preprocessing, evaluation or idle provisioned time: USD 4.26 projected against USD 29.76 realised on the variant-effect study. Neither error is a property of gating as such; both are properties of what we chose to meter, and both are cheap to correct once measured rather than assumed.

### How far each result reaches

On host range the accuracy comparison against the general agent is a tie and we report it as one: 0.826 against 0.812 without confidence intervals is not a win, and what the run does separate is process cost and reliability. On hit-to-lead the gain decomposes into two interpretable mechanisms rather than one opaque fit, since a monotonic calibration cannot reorder anything and the ranking gain therefore has to come from the QSAR blend; its boundaries, an empty pre-registered contrast and a physics anchor that mapped nothing, were surfaced by the system itself. On variant effect the adapted model is better calibrated than either baseline without being well calibrated, the region-novel interval contains zero so we do not claim generalisation to unseen protein regions, and the 1,000 Cohort C variants are prioritisation candidates rather than reclassifications.

### Execution success is not scientific correctness

This paper contains two instances of that failure mode, and only one is somebody else’s. The general coding agent produced read-window fragments scoring 0.00% against a blinded golden set while running to completion without error; CAPO in turn shipped a preprocessing regression in the variant-effect study that passed its own runtime checks and was caught only by the leakage audit. In neither case was execution status informative about scientific validity, and what caught both were domain checks outside the generating system: an independently curated golden set, a temporal split, an explicit leakage audit. Generated data transformations should therefore be treated as scientific claims and tested as such, by default and not on suspicion. For a field moving toward closed loops this is the more consequential of our two findings: an agent that spends too much is an accounting problem and the loop merely runs slower; one that silently corrupts its inputs at the front of the pipeline, where a naive evaluation never looks, invalidates everything downstream while reporting success.

### Limitations

Each task is a single end-to-end run, without seeds or variance estimates on the agentic process itself, and the gate is evaluated only as a whole, so zero-versus-four failed trials is a natural experiment and not an ablation. The head-to-head covers one task; we ran it on host-range prediction because that is where it is most informative, the only task with a hand-designed expert architecture as an independent reference and by far the largest labelled corpus. We expect the pattern to hold wherever raw data must be prepared and a budget binds, but one task cannot establish it. Each study carries the bounds reported above: the empty pre-registered FEP+ contrast, the two-target Boltz-hard case, the unmapped physics anchor, the small pathogenic-skewed variant panel, an ECE of 0.190, a region-novel interval including zero. There is no closed loop and no wet-lab validation, and the scope is a prototype: three tasks, two backbone families, a curated registry, and protocols that needed biological expertise to design.

### Conclusion

Two assumptions fail once the tool is a fine-tuning run: that it is cheap and reliable, and that a run which completes is right. We reframed adaptation as a gated decision problem to deny both: a probe measured on the target accelerator, a budget gate allowed to decline, a canary inside the real training loop, an audit trail in which a missing artifact is a hard failure. One unchanged system under an identical USD 50 budget then chose three fit-for-purpose techniques, a refusal among them; beat a hand-designed expert transformer; matched a general coding agent’s accuracy with 57.4% less GPU time, 32.5% less cost per successful run and no failed trials; and recovered the target domains at 97.6% and 99.83% where that agent silently returned unusable output. On this evidence, adaptation can be made systematic, auditable and compute-aware. CAPO does not supply the judgement that defines and validates a biological task; it connects that judgement to the compute needed to act on it, which could bring adaptation within reach of laboratories that generate assay data, not pipelines. The components mostly exist; the next steps follow from our limitations: whole-run budget control, golden-set tests on generated preprocessing, ablations tight enough that a refusal is decisive. A closed-loop laboratory that proposes its own experiments will sit on a layer deciding what to compute and what to trust; the contribution here is that layer, and the finding that its most valuable decision may be a refusal.

## A Cross-task synthesis

Table 4 places the three runs of Section 5 side by side. It is a summary of results reported in full in the body, not a separate result.

**Table 4:** One unchanged system, three fit-for-purpose adaptation techniques. Realised costs are not comparable across rows (Section 6); host-range totals are read from Figure 5.

| Task | Data regime | Gate decision | Headline result | Realised cost |
| --- | --- | --- | --- | --- |
| Host range | Raw NGS reads → 4.90M dev / 0.44M test examples, imbalanced | **Admit:** partial fine-tune, ESM-2 8M, last four layers unfrozen | Macro-MCC 0.826 vs 0.743 expert; matched vs general agent; 0 vs 4 failed trials | 3h 52 GPU, USD 17.09 total |
| Hit-to-lead | 644 cached co-folds; 15 scorable series; sparse labels | **Decline:** freeze Boltz-2, fit assay-calibrated head on outputs | Spearman 0.700 vs 0.594; better on 11/15 targets; pre-registered contrast empty | 1.68 GPU-h, USD 3.35 compute |
| Variant effect | 1,882 training labels, pathogenic-skewed; temporal no-lookahead split | **Admit:** rank-4 LoRA on ESM-2 150M, three seeds | Temporal MCC 0.478 vs 0.252 / 0.000; region-novel interval includes zero | ~1.5 h, USD 29.76 (4.26 projected) |

## B Adaptation prior and evaluation protocol

Table 5 gives the deliberately conservative default adaptation strategy, keyed on label count, from which every run is initialised. It is a starting point, not a rule: the final choice depends on task type, label structure and quality, sequence lengths, backbone size, budget, overfitting risk and the expected quality of frozen embeddings, and is settled only by the Phase 3 gates. Two of the three studies reported here ended on a strategy other than the one the prior nominated. The planned epoch count *E* is resolved in priority order, an explicit user configuration superseding a model-card recommendation surfaced by the Hugging Face researcher, which in turn supersedes a family default.

**Table 5:** Default adaptation prior, keyed on label count.

| Label count | Default adaptation strategy |
| --- | --- |
| 0 | Zero-shot inference (no training) |
| < 500 | Linear probe over a frozen backbone |
| 500 to 10k | Parameter-efficient tuning, LoRA with $r = 8$ to 16 |
| 10k to 100k | Higher-capacity LoRA, $r = 16$ to 32 |
| > 100k | Full fine-tuning, warm-started from a LoRA checkpoint |

Table 6 gives the post-training evaluation protocol. Evaluation is always on data excluded from both optimisation and online model selection.

**Table 6:** Post-training evaluation protocol.

| Component | Protocol |
| --- | --- |
| Test data | An existing held-out test set is preserved; otherwise CAPO constructs a new split. |
| Split strategy | Sequence-aware where appropriate: homologous sequences are clustered and assigned at cluster level, so generalisation is measured beyond near-duplicates. |
| Classification | AUROC, AUPRC, accuracy, MCC and calibration error as appropriate; AUPRC and MCC prioritised under imbalance. |
| Regression | RMSE, MAE, Pearson and Spearman, selected according to whether the task emphasises absolute accuracy or rank ordering. |
| Baselines | The adapted model is evaluated on the same partitions as frozen-representation models, standard adaptation methods and user-provided expert baselines. |
| Robustness | Where external or shifted datasets exist, CAPO tests whether the improvement persists beyond the primary test distribution. |

## C Full agent roster

Table 7 lists the eleven agents, the phase each runs in, the model it is pinned to and the role that allocation is matched to.

**Table 7:** CAPO agents, with the role each model allocation is matched to. Model capacity is matched to the complexity, frequency and scientific risk of each role; tool permissions follow the same principle and are fixed in code, so every agent has an explicit, inspectable capability surface.

| Agent | Phase | Model | Role |
| --- | --- | --- | --- |
| Hugging Face researcher | Pre-run | Sonnet 4.6 | Bounded retrieval of external evidence under a strict output contract. |
| Memory consultant | Pre-run | Haiku 4.5 | Retrieval of relevant evidence from previous CAPO runs. |
| Infrastructure | 0 | Opus 4.8 | Budget-aware environment selection from live pricing and available capacity. |
| Data profiler | 1 | Opus 4.8 | Interpretation of heterogeneous inputs and definition of preprocessing decisions. |
| Model selector | 2 | Sonnet 4.6 | Rubric-based ranking of candidates from a curated registry. |
| Orchestrator | 3–4 | Opus 4.8 | Open-ended planning, code synthesis, probe and cost reasoning, failure recovery. |
| Experiment tracker | 4 | Haiku 4.5 | Initialisation of the experiment-tracking record. |
| Health monitor | 5 | Haiku 4.5 | Repeated read-only classification of training health. |
| Recovery agent | Recovery | Opus 4.8 | Root-cause diagnosis of post-canary failures and bounded repair. |
| Code-repair critic | Repair | Opus 4.8 | Diagnosis and repair of failures unresolved after standard correction attempts. |
| Finaliser | 6 | Opus 4.8 | Diagnosis of the terminal state and evidence-based report generation. |

**Table 8:** Blinded evaluation of raw-read processing on two host libraries not used for modelling. The golden set was produced by a reference assay pipeline and withheld from both systems during preprocessing; the match rate is the proportion of recovered sequences agreeing with it.

| System | Output | Match | Cost |
| --- | --- | --- | --- |
| *Pangolin host* |  |  |  |
| CAPO | full-length canonical RBDs (14.4M) | **97.6%** | \$9.91 |
| General agent | read-window fragments | 0.00% | \$25.96 |
| *Possum host* |  |  |  |
| CAPO | canonical 201-aa RBDs (2.61M) | **99.83%** | \$6.88 |
| General agent | short fragments (79 to 167 aa) | 0.00% | \$7.09 |

**Table 9:** Deterministic construction funnel for the variant-effect task, from the attrition log. More than two million candidate expressions are filtered to a small, quality-controlled set of labelled variants and three locked cohorts.

| Construction step | Retained | Note |
| --- | --- | --- |
| Panel rows, historical release | 36,887 | GRCh38, eight genes |
| Panel rows, current release | 85,371 | endpoint release |
| Simple missense HGVS expressions | 2,285,385 | across the panel |
| Matching MANE Select accession | 69,914 | wrong-transcript expressions dropped |
| Historical missense, residue-validated | 16,977 | one residue mismatch dropped |
| Historical definitive labels | 2,897 | 2,451 pathogenic, 446 benign |
| Cohort A, development set | 1,882 | train 1,312 / val 279 / test 291 |
| Cohort B, temporal reclassified VUS | 420 | 198 to pathogenic, 222 to benign |
| Cohort C, currently uncertain VUS | 1,000 | prospective prioritisation only |

## D Host-range task: data, curation and the expert baseline

The data come from a yeast-surface-display screen of a combinatorial mutagenesis library of the Omicron BA.1 RBD, residues 331 to 531 (201 residues), stained against a panel of animal ACE2 orthologs and sorted by fluorescence-activated cell sorting into binding and non-binding populations before deep sequencing [25] (Figure 6). CAPO ingests the raw sequencing output together with assay metadata specifying the target region, the flanking structure, the reading frame and the label-construction rule. Its data profiler translates reads to amino-acid sequences, strips primers and flanks, deduplicates and assigns a binder or non-binder label per ortholog, yielding roughly 9.5 million labelled examples at 201 residues (Figure 7). Ten orthologs have sufficient single-label support: human, mouse, rat, dog, cat, cattle, horse, bat, monkey and mink. Restricting to examples with exactly one observed species label leaves 4,895,620 examples for development and 441,865 for held-out testing, with a further tenth of the training data reserved for validation and threshold tuning. The dataset ships precomputed homology-aware sequence clusters (one-hot encoding, incremental PCA, *k*-means into five clusters) that CAPO reuses to keep related variants in the same partition. Binder prevalence ranges from 3.2% (rat) to 51.4% (mink) with a cross-species mean of 21.4% (Figure 8).

**Figure 4:**
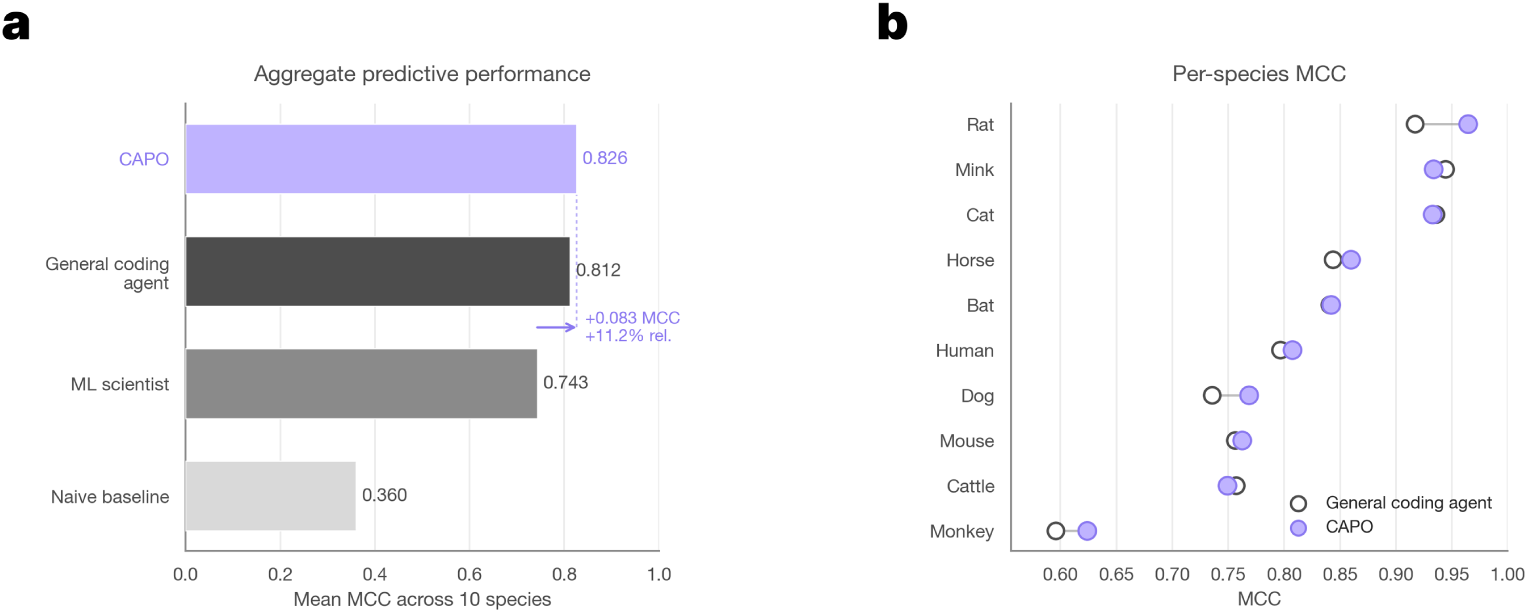
Host-range predictive performance, both panels. (a) Species-averaged MCC per method. (b) Per-species MCC under tuned decision thresholds; CAPO matches or exceeds the general coding agent on 7 of the 10 orthologs, with gains concentrated on the lower-MCC species. Panel (a) is reproduced in Figure 2.

**Figure 5:**
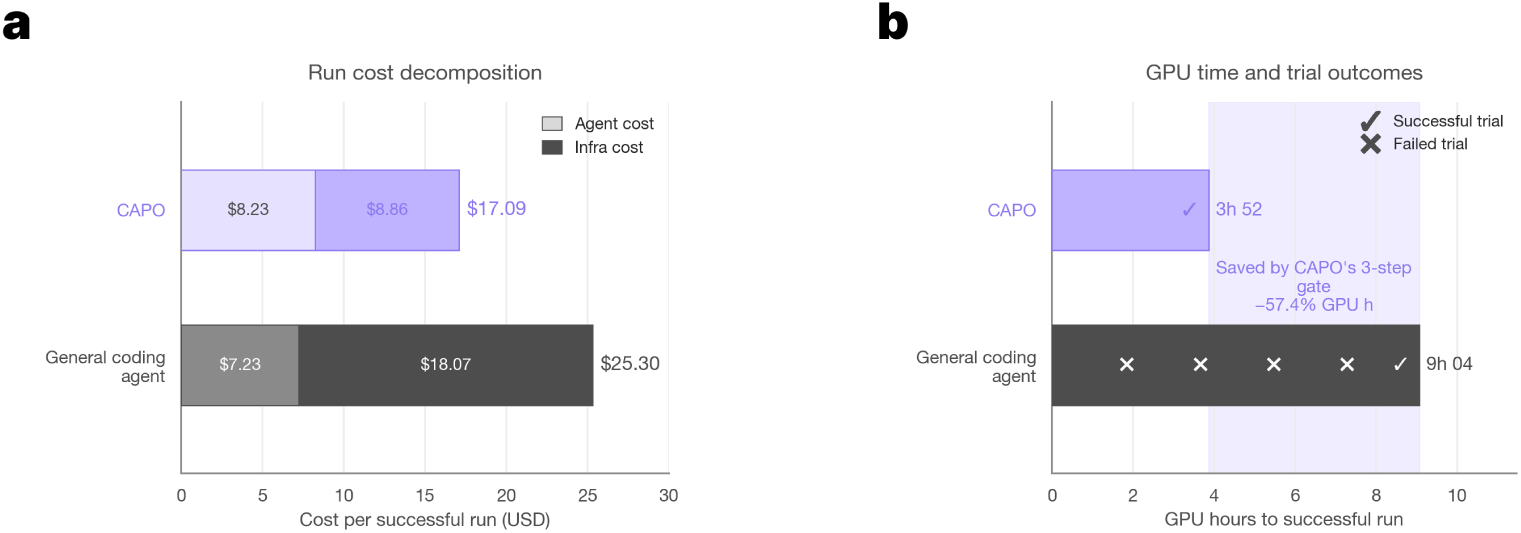
Host-range process cost, both panels. (a) Cost per successful run decomposed into agent and infrastructure components. (b) GPU hours to a successful run, with failed trials marked; reproduced in Figure 2. The per-successful-run totals of USD 17.09 (CAPO) and USD 25.30 (general agent) quoted in Table 4 are the printed labels of panel (a).

**Figure 6:**
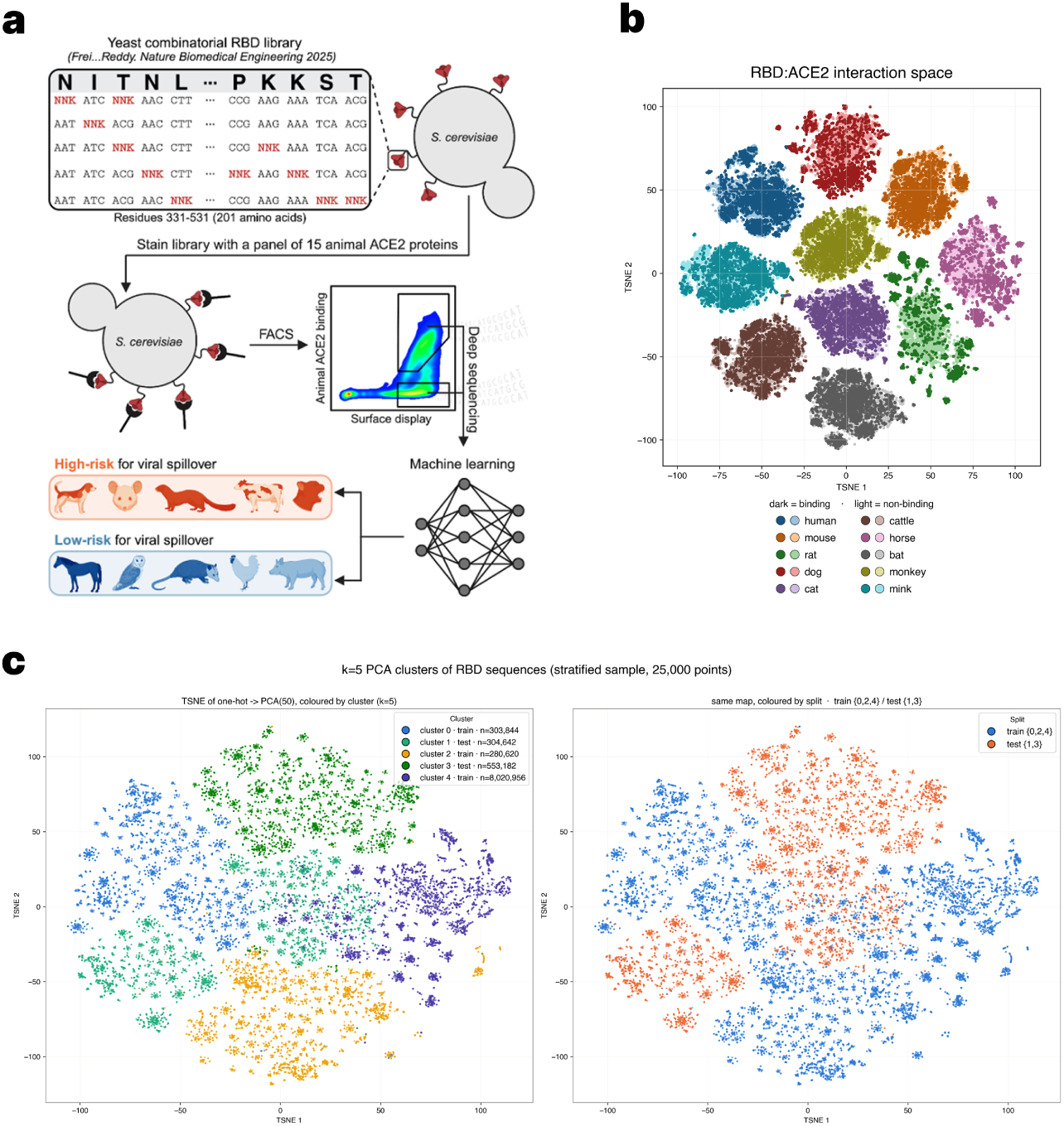
The host-range platform. (a) The combinatorial yeast-display library is stained against a panel of animal ACE2 orthologs, sorted by fluorescence-activated cell sorting, deep sequenced and used to train a model that flags high-risk and low-risk hosts. (b) An embedding of the RBD:ACE2 interaction space, coloured by ortholog and by binding outcome. (c) An embedding of RBD sequence space coloured by sequence cluster and by a candidate cluster split, showing the homology structure available for leakage-controlled evaluation.

**Figure 7:**
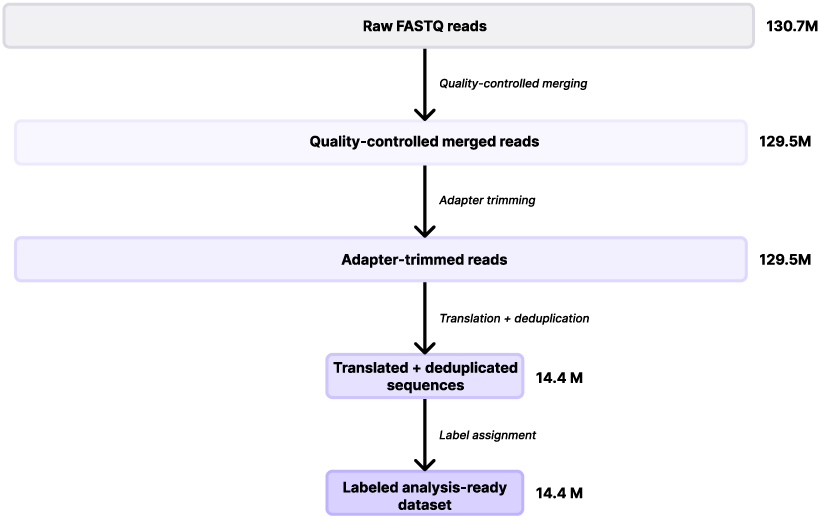
From raw reads to model-ready sequences. Quality control, trimming, filtering and deduplication remove technical noise while preserving biological signal: low-quality reads and variants observed fewer than twice are discarded and survivors are translated to amino acids, yielding multi-label training sets annotating each variant across species.

**Figure 8:**
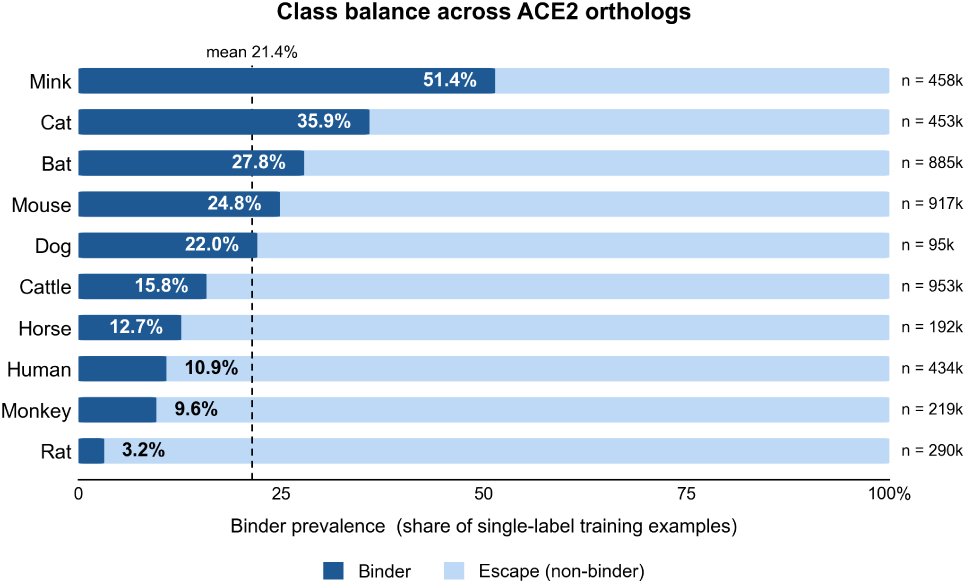
Class balance across the ten ACE2 orthologs in the training set. Bars give the share of single-label training examples labelled as binders, with per-species counts on the right and the cross-species mean of 21.4% marked.

Because the assay reports binding or non-binding to exactly one ortholog per retained sequence, with the other nine unknown, CAPO formulates the task as multi-task binary classification with partially observed labels: one model reads the RBD sequence and emits ten binding logits, only the observed label contributes to the loss and the remaining outputs are masked. This avoids treating unmeasured interactions as non-binding while letting the shared backbone learn from all species. ACE2 sequences are not supplied as inputs, so predictions rest on the RBD sequence alone, and the evaluation therefore demonstrates generalisation to unseen RBD variants across the ten modelled orthologs, not to receptors from species absent during training.

### How the blinded curation score is computed

The host-range task opens with a data-engineering problem that is easy to underestimate: the raw material is paired-end NGS output from a mutagenesis library, not a table of sequences, so turning it into supervised examples means merging read pairs, trimming adapters and flanks, filtering low-quality and rare reads, and translating each survivor into its canonical receptor-binding domain. We score that step blind, against an independently curated golden set of full-length RBD sequences withheld from both systems while they preprocess. The golden set was produced by a reference assay pipeline that quality-filters and merges the raw reads, trims the library adapters to isolate the variable region, reattaches the constant flanks, and translates each read into its full-length 201-residue domain. The match rate we report is the proportion of the sequences a system recovers that agree with the corresponding golden-set sequence.

To separate preprocessing quality from downstream prediction performance, sequence recovery was evaluated on two host libraries, pangolin and possum, that were not used in the modelling study. The same assay-processing procedure was applied to both with no species-specific tuning, so they are independent tests of whether that procedure reconstructs full-length domains beyond the species it was exercised on. CAPO agreed with the golden set on 97.6% of the 14.4M pangolin sequences and on 99.83% of the 2.61M possum sequences (Table 8). The residual few per cent on pangolin are sequences CAPO recovered that the reference pipeline resolved differently; they are reported as disagreements rather than adjudicated, because the golden set is the reference by construction here.

### Why the general agent’s score is exactly zero

The criterion is deliberately strict: agreement means the recovered sequence is exactly the golden-set sequence, and the golden set contains fulllength canonical domains. Given identical raw reads, the general coding agent stopped after merging read pairs and emitted read-window fragments of 79 to 167 residues, which cannot span a 201-residue domain, so no fragment can match and the rate is 0.00% rather than merely low. We state this plainly because it changes how the number should be read. It is not a near miss under a harsh metric, and a partial-credit criterion would change the number without changing the conclusion: a fragment that omits part of the domain also omits any mutations falling outside it, so distinct library variants collapse onto identical fragments and the per-variant binding labels can no longer be attached to the variant they describe. The dataset the task asks for does not exist in that output. The failure is equally invisible from inside the run, since both agent runs terminated normally and returned plausibly formatted sequence files, which is exactly why the step is scored against a reference the generating system never sees.

### The expert baseline

The strongest human comparator (Figure 9) is a per-residue transformer that scores one RBD:ACE2 pair at a time. Per-residue interaction features pass through a linear projection with learned positional embeddings into a transformer encoder with padding masks over the padded RBD positions. The encoder output passes through a position-wise feedforward network that aggregates residue-level embeddings into a compact lower-dimensional representation, which is flattened across positions and fed to a linear head emitting one logit per RBD:ACE2 pair, trained under binary cross-entropy. It is a carefully engineered model refined over multiple iterations for this specific assay, which is what makes it the right target for a system that never writes a line of architecture code.

**Figure 9:**
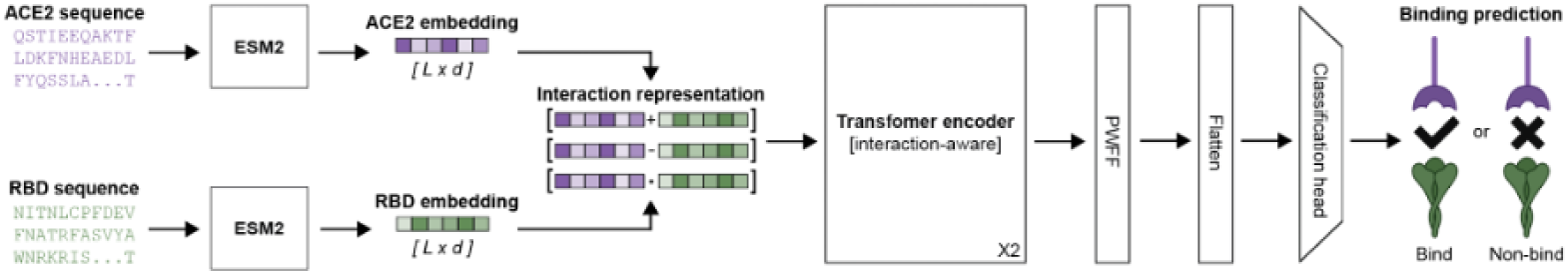
The expert baseline architecture: a hand-designed per-residue transformer predicting binding for one RBD–ACE2 pair at a time.

## E Hit-to-lead task: data, methods and extended results

Evaluation uses the public FEP+ binding free-energy benchmark, a standard medicinal-chemistry set of congeneric compound series with measured binding free energies across pharmaceutically relevant targets [26, 27]. CAPO co-folded 644 protein–ligand complexes across the benchmark’s 29 targets with Boltz-2, reusing co-folds computed on a previous run so that no accelerator time was spent recomputing them. Each measured Δ*G* in kcal mol*^−^*^1^ becomes a potency label log_10_(*K_d_/µ*M) = Δ*G/*1.364 + 6, lower being stronger. Targets whose held-out batch contains fewer than three compounds are not scored individually, leaving 15 targets with a stable per-target ranking metric. Compound names are parsed into a series prefix and a numeric suffix, the earliest 65% of each series forming the measured first batch and the remainder the held-out second batch, with a Murcko-scaffold fallback when names lack a usable ordering. Second-batch labels are touched exactly once, at final scoring.

Four methods are compared on identical co-folds and splits (Figure 10). Raw Boltz-2 ranks by its affinity score directly. Calibrated Boltz-2 applies a per-target linear map to the potency scale, a monotonic transform that corrects magnitude without changing rank order. The pooled classical control is a single kernel-ridge regressor with an RBF kernel, fitted once over the pooled first batches of all targets on physicochemical descriptors and frozen Boltz-2 scalars, with identical hyperparameters and no per-target adaptation. The CAPO head calibrates Boltz-2, assembles features from Boltz-2 signals, interface contacts read from the predicted complex, extended-connectivity fingerprints [31] and physicochemical descriptors, fits a leave-one-out-gated ensemble of ridge and random-forest regressors, and blends it with the calibrated Boltz-2 signal by a weight learned on the first batch alone. Boltz-2’s weights are never updated in any of the four, which is what makes reusing cached co-folds valid.

**Figure 10:**
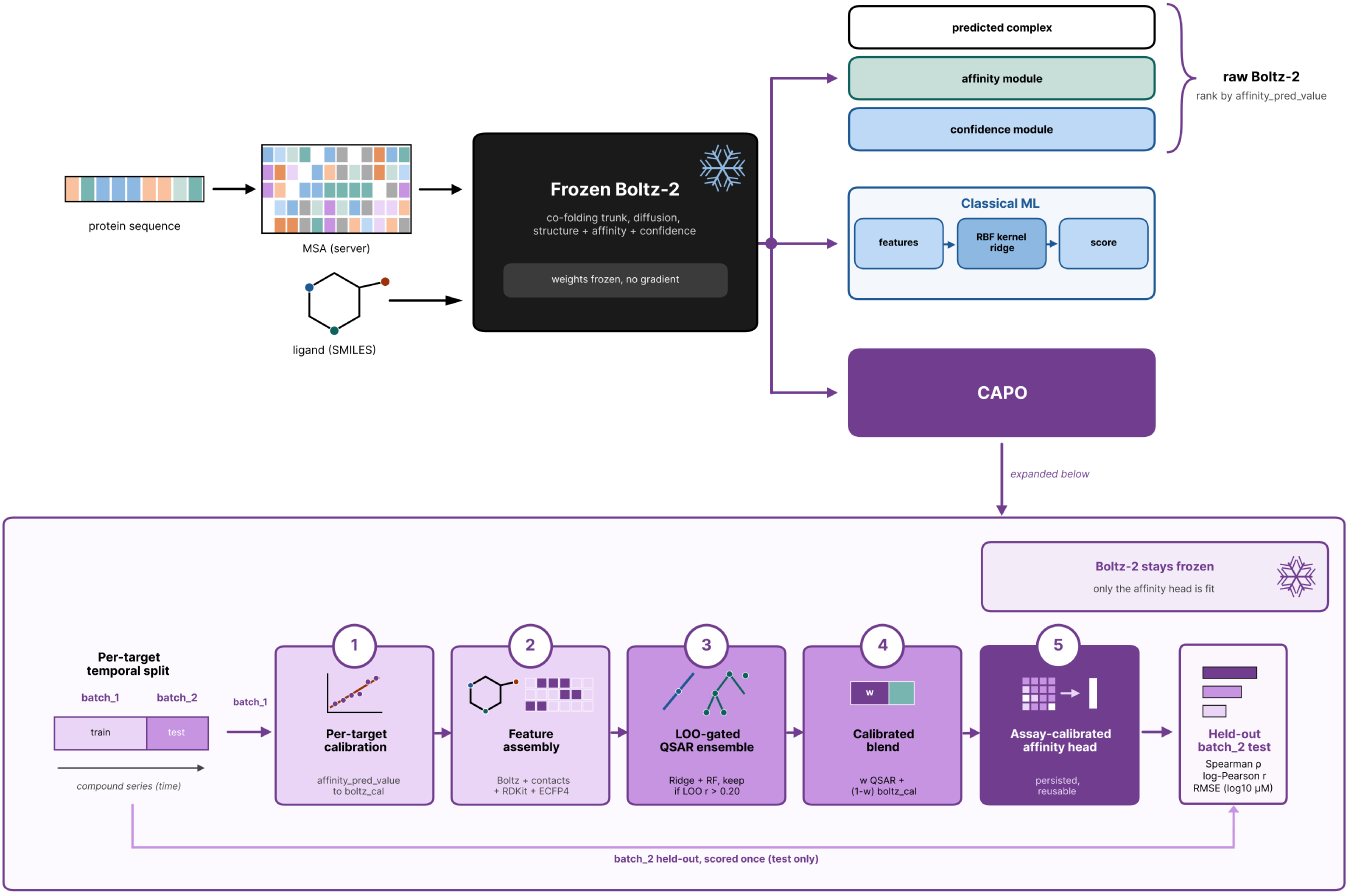
The hit-to-lead pipeline. Protein sequence and ligand SMILES are co-folded by frozen Boltz-2, whose structure, affinity and confidence outputs feed three comparators. The CAPO head uses a per-target temporal split, calibrates the frozen affinity score, assembles Boltz-2, contact and chemical features, fits a leave-one-out-gated QSAR ensemble, blends it with the calibrated signal, and is scored once on the held-out batch.

## F Variant-effect task: construction and extended results

The eight-gene hereditary-cancer panel is BRCA1, BRCA2, ATM, PALB2, TP53, PTEN, MLH1 and MSH2, spanning homologous-recombination and DNA-repair genes, mismatch-repair genes and tumour-suppressor signalling genes. CAPO links the historical and current ClinVar summaries to a protein-level HGVS map, retains GRCh38 variants from the eight genes, and keeps only simple missense substitutions affecting a single amino acid. Each variant is mapped to the gene’s MANE Select protein [32], and CAPO verifies that the reference amino acid in the HGVS annotation matches the residue at the reported position; variants with inconsistent transcript mappings or residue numbering are removed before they can enter as silently mislabelled examples. Labels use only the aggregate germline classification with an accepted review status, collapsed to pathogenic or benign, excluding conflicting, risk-allele, drug-response and other ambiguous categories. Table 9 traces the construction funnel. Figure 14 sketches the temporal design.

**Figure 11:**
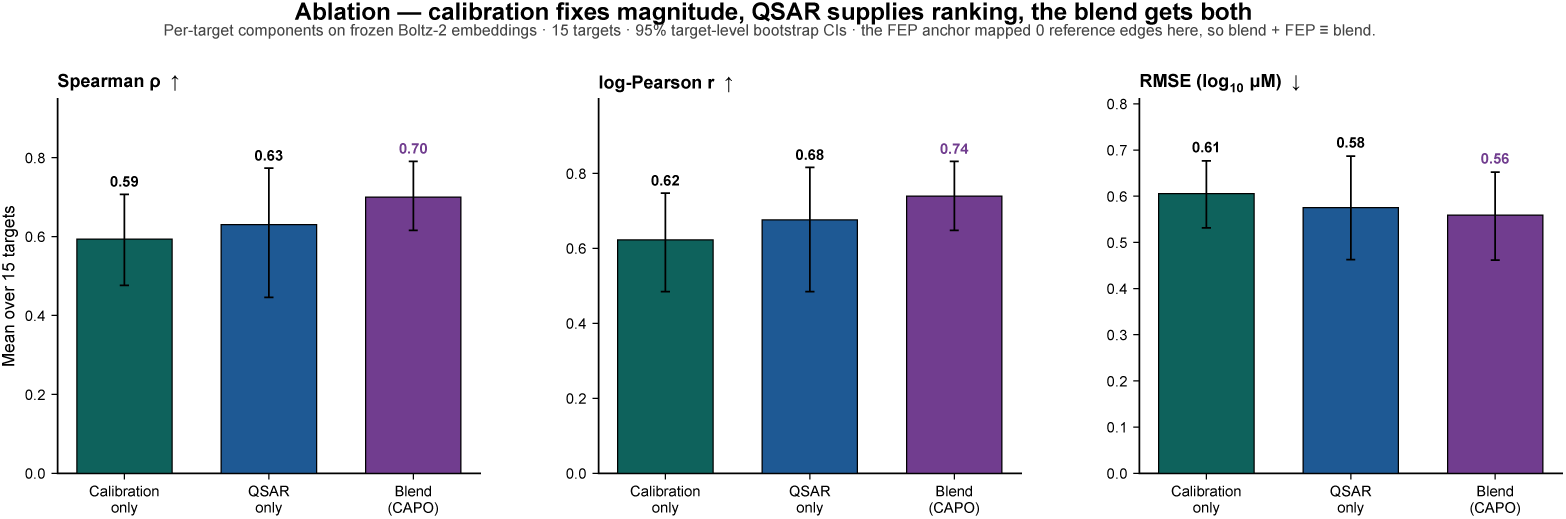
Ablation decomposition. The calibration is a monotonic map and cannot reorder anything, so the whole ranking gain is contributed by the QSAR blend, while calibration does almost all of the absolute-error reduction.

**Figure 12:**
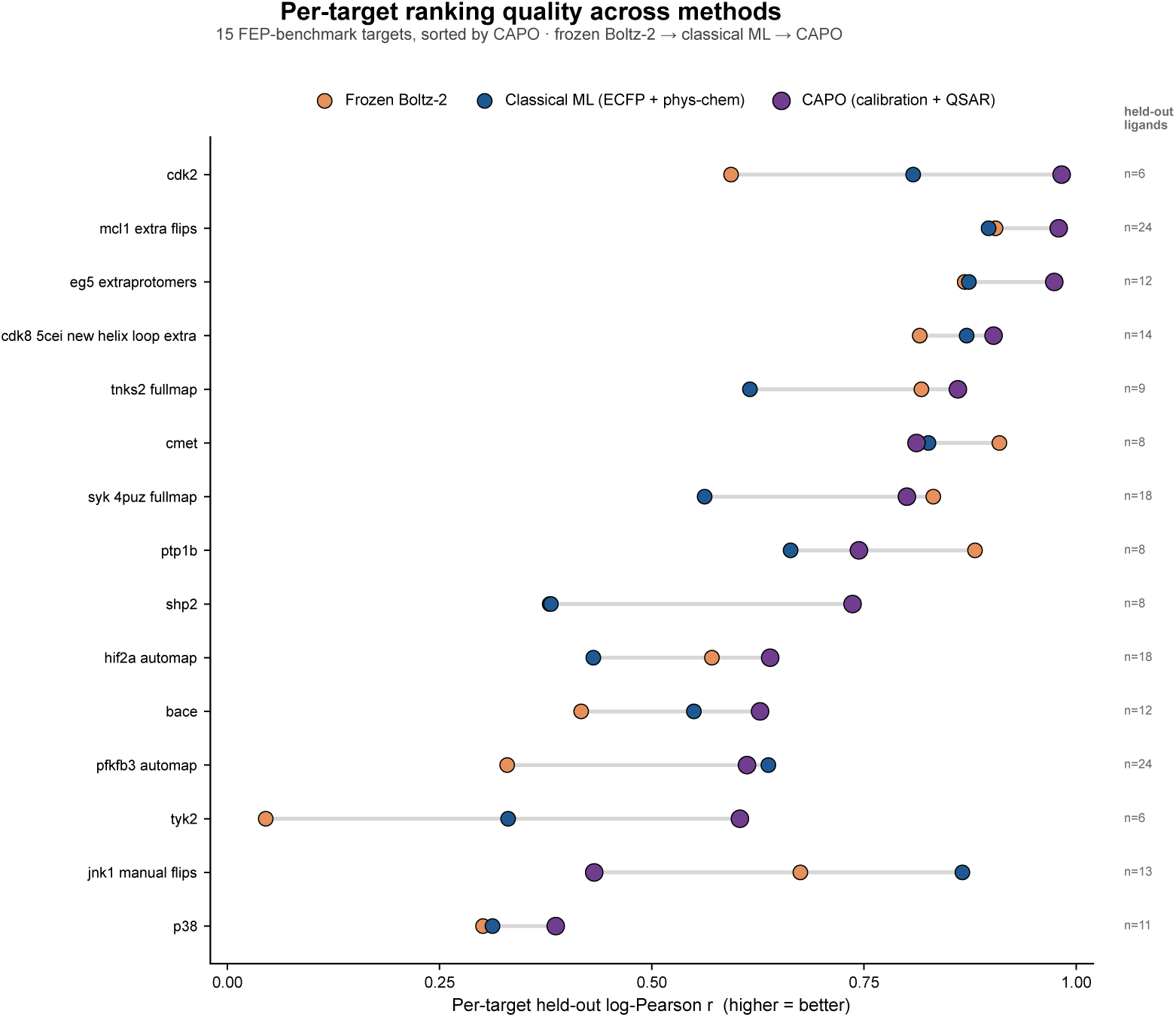
Per-target breadth in full, including the targets omitted from Figure 3.

**Figure 13:**
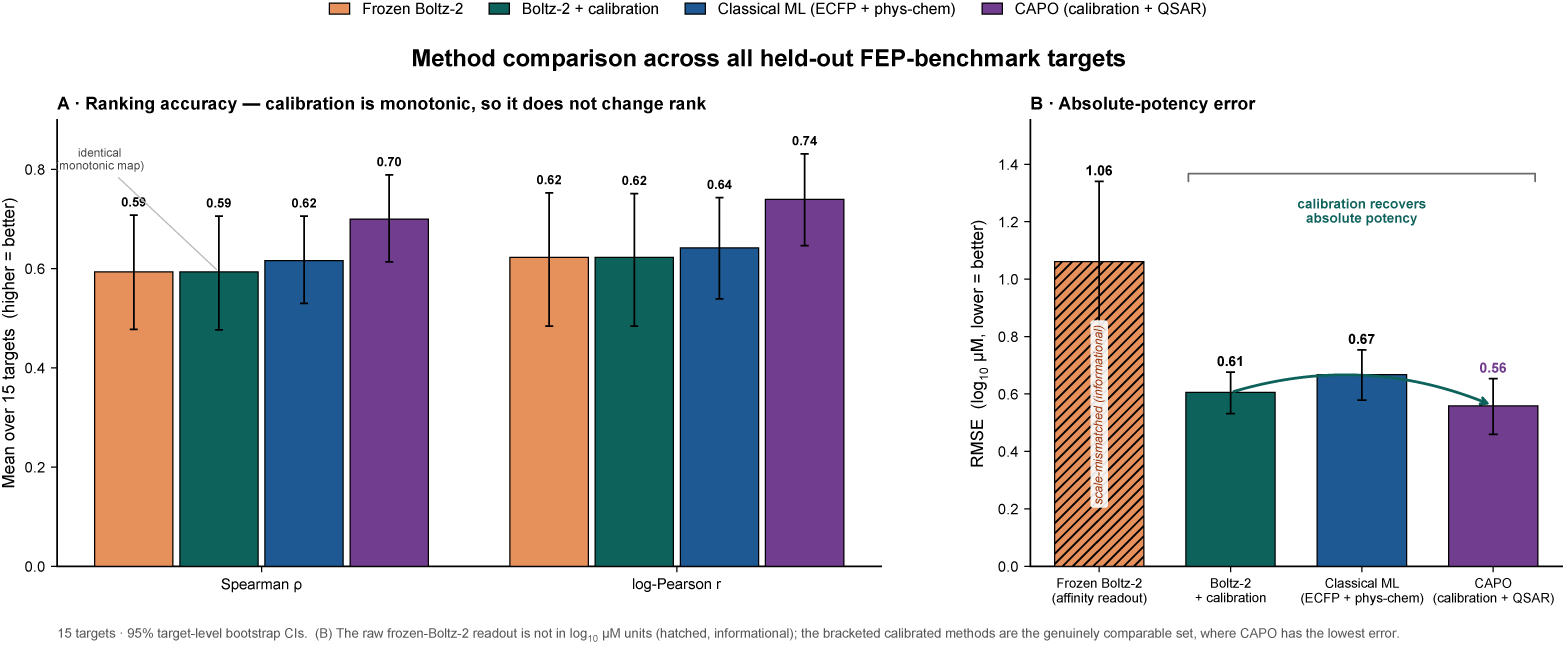
Method comparison across the scored FEP+ series.

**Figure 14:**
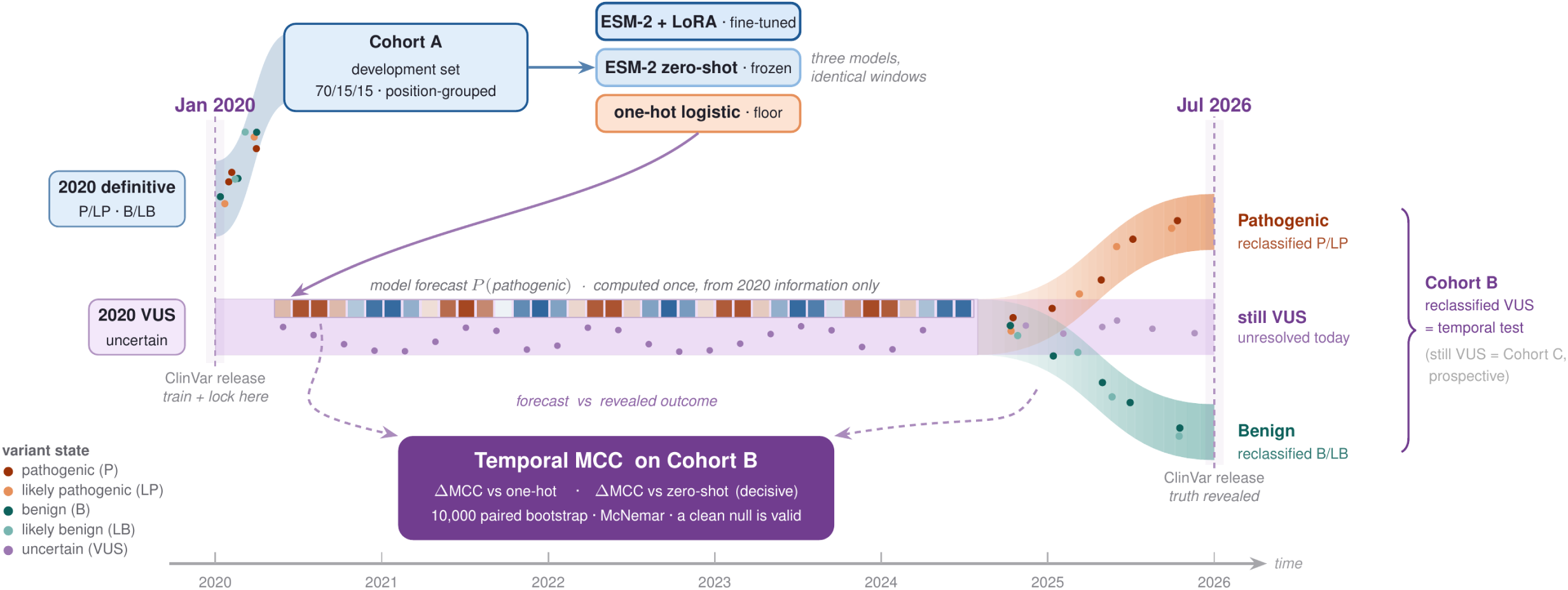
Temporal no-lookahead design of the variant-effect benchmark. All three models are fitted and thresholded on Cohort A, the variants already classified pathogenic or benign in the January 2020 ClinVar release, and each produces one forecast from 2020 information alone. Cohort B, the primary endpoint, consists of variants that were uncertain in 2020 and had been reclassified by the July 2026 release; variants still uncertain today form Cohort C, which is scored for prospective prioritisation only.

Cohort A is capped and stratified by gene and label so that no gene dominates a class, then split with a grouped split that keeps every substitution at the same gene and residue position in the same partition. Cohort B, the primary endpoint, was held completely hidden until the models and their decision thresholds were locked. Cohort C is scored for prospective prioritisation only and never treated as ground truth. The one-hot baseline encodes both windows and concatenates the reference, the alternate and their difference into a logistic regression. The field-standard control is the pretrained ESM-2 150M [1] zero-shot masked-marginal log-likelihood ratio at the mutated position [33]. The adapted model applies rank-4 low-rank adaptation [30] to the same backbone, passing reference and alternate windows through a shared encoder and forming a mutation representation from the two residue embeddings, their difference and its magnitude before a small classification head; only the adapters and the head are trained, across three seeds whose predictions are averaged.

## G Projected versus realised cost

CAPO projects a cost before it launches and reconciles it against measured wall-clock cost at finalisation. Across the three studies the gate overestimates realised GPU cost by roughly 2.1*×* to 3.2*×* (Figure 18), reserving headroom that keeps each study inside its budget once agent cost is added. Overestimating is the direction we prefer, because it means the budget is not crossed, but it is a real limitation: a gate that is systematically 3*×* pessimistic will decline configurations that would in fact have fitted.

**Figure 15:**
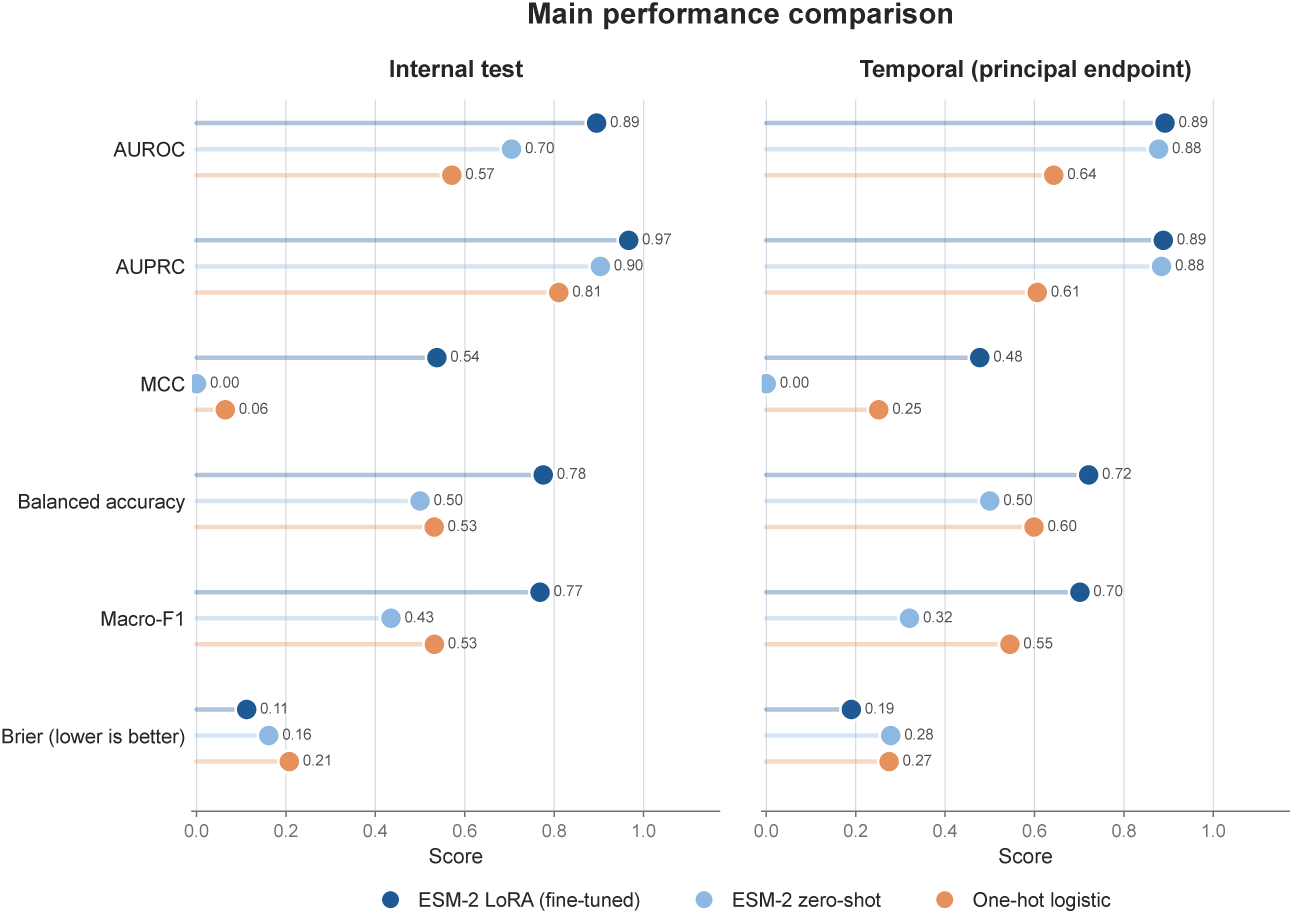
Performance with 95% bootstrap intervals for all three models on the internal test set (left) and the temporal reclassified-VUS benchmark (right, the principal result). The adapted model dominates the one-hot floor on every metric and clears the frozen zero-shot control on threshold-dependent metrics such as MCC and balanced accuracy.

**Figure 16:**
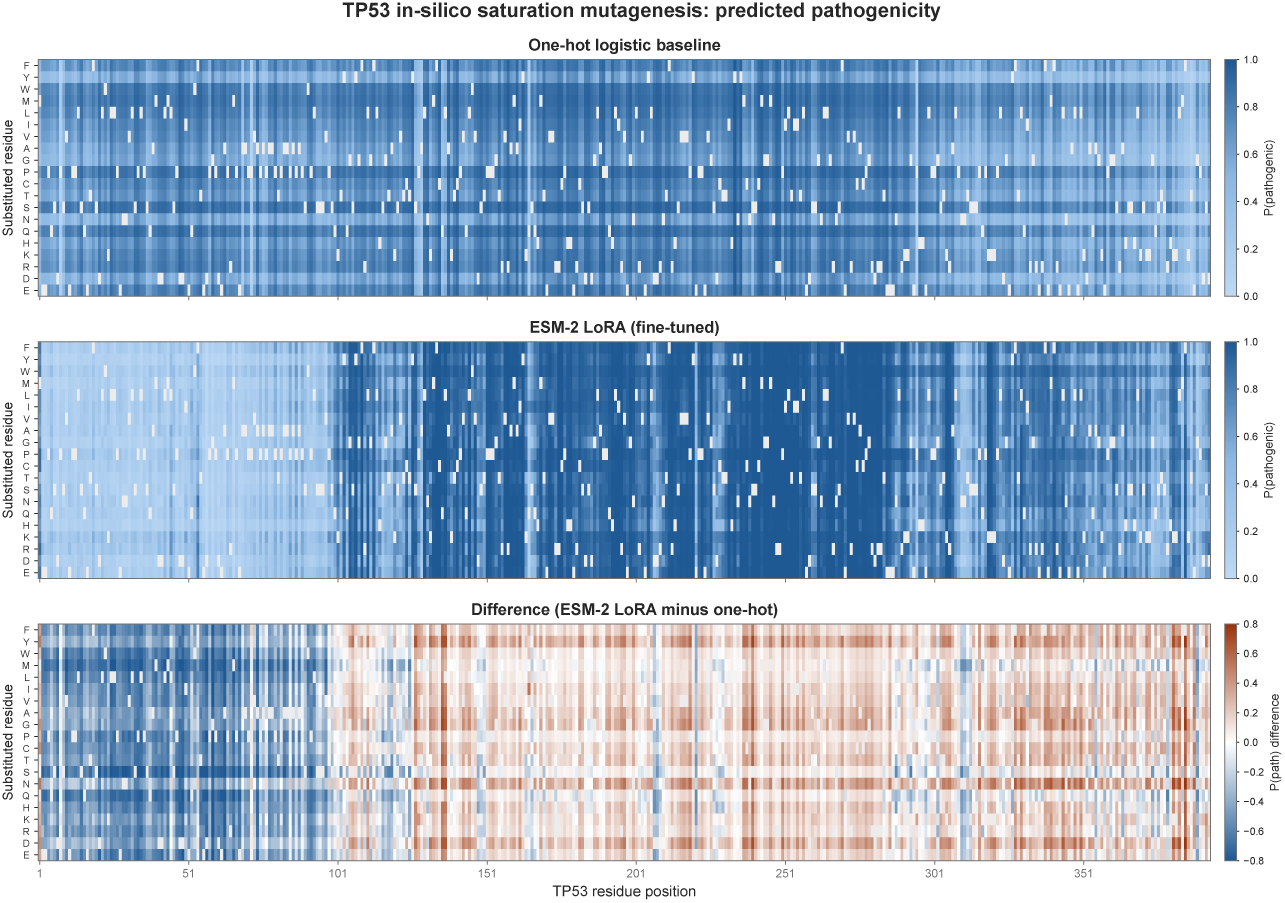
In-silico saturation mutagenesis of TP53, scoring all 393 *×* 19 single substitutions. Top: one-hot baseline pathogenicity. Middle: adapted ESM-2. Bottom: their difference. The adapted model recovers coherent, position-specific mutation-sensitive regions over the DNA-binding core that the diffuse baseline misses. This map is illustrative and is not validated against deep mutational scanning.

**Figure 17:**
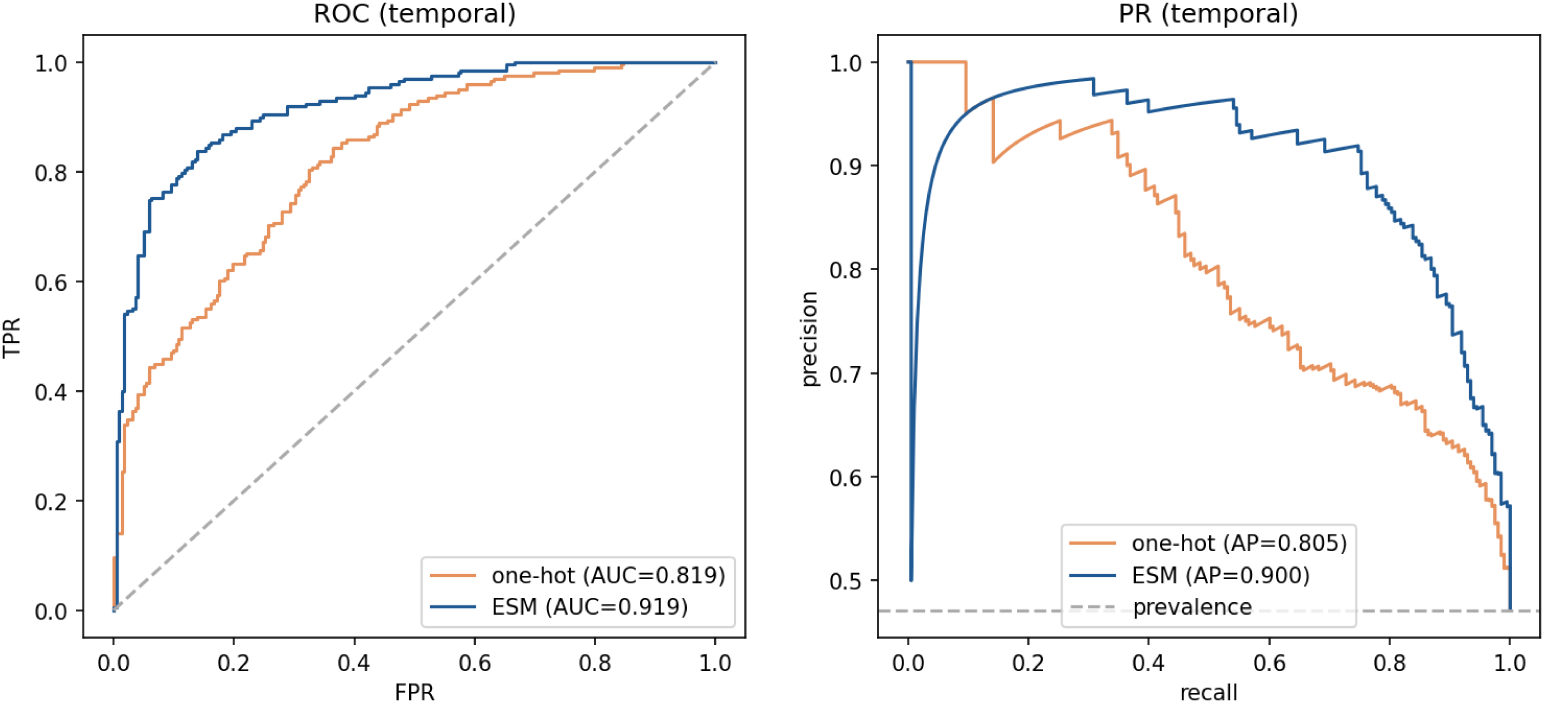
ROC and precision–recall curves on the temporal cohort.

**Figure 18:**
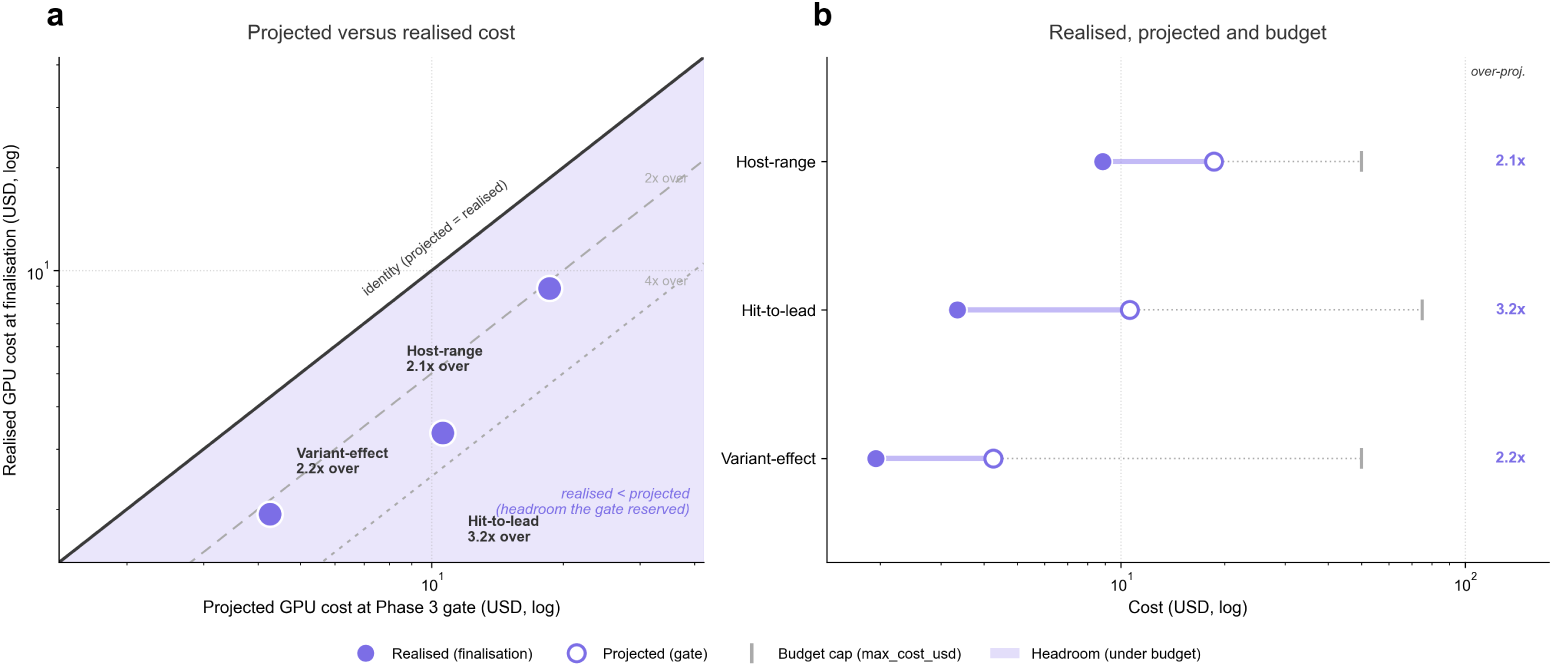
Projected against realised cost for the three runs. (a) Realised GPU cost at finalisation against the projected GPU cost from the Phase 3 gate, with the diagonal marking a perfect projection. (b) The same runs as realised, projected and budget-cap triples on a common axis.

## H Agent model pricing

Table 10 reports the prices used to compute the agent costs throughout, as of 16 July 2026. The mechanism described in Section 3 is model-agnostic; the specific allocations and prices are not.

**Table 10:** Claude API pricing used for cost reporting, as of 16 July 2026, in USD per million tokens. Prompt caching is used extensively during runs and reduces the cost of repeatedly supplying shared instructions and context.

| Model | Input | 5-min cache write | 1-h cache write | Cache hit | Output |
| --- | --- | --- | --- | --- | --- |
| Claude Opus 4.8 | \$5.00 | \$6.25 | \$10.00 | \$0.50 | \$25.00 |
| Claude Sonnet 4.6 | \$3.00 | \$3.75 | \$6.00 | \$0.30 | \$15.00 |
| Claude Haiku 4.5 | \$1.00 | \$1.25 | \$2.00 | \$0.10 | \$5.00 |

## I Example run report

Every run is finalised into a report in which each claim derives from a specific artifact and any unavailable value is recorded as unknown. The report below is reproduced verbatim from a host-range run, with dataset and repository identifiers replaced by placeholders. It is a different run from the one reported in Section 5.1, included here to show the artifact format in full, so its headline figures (macro-MCC 0.8229, USD 38.43 total, 6h 09m instance runtime) are its own and not those of Figures 2 and 5.

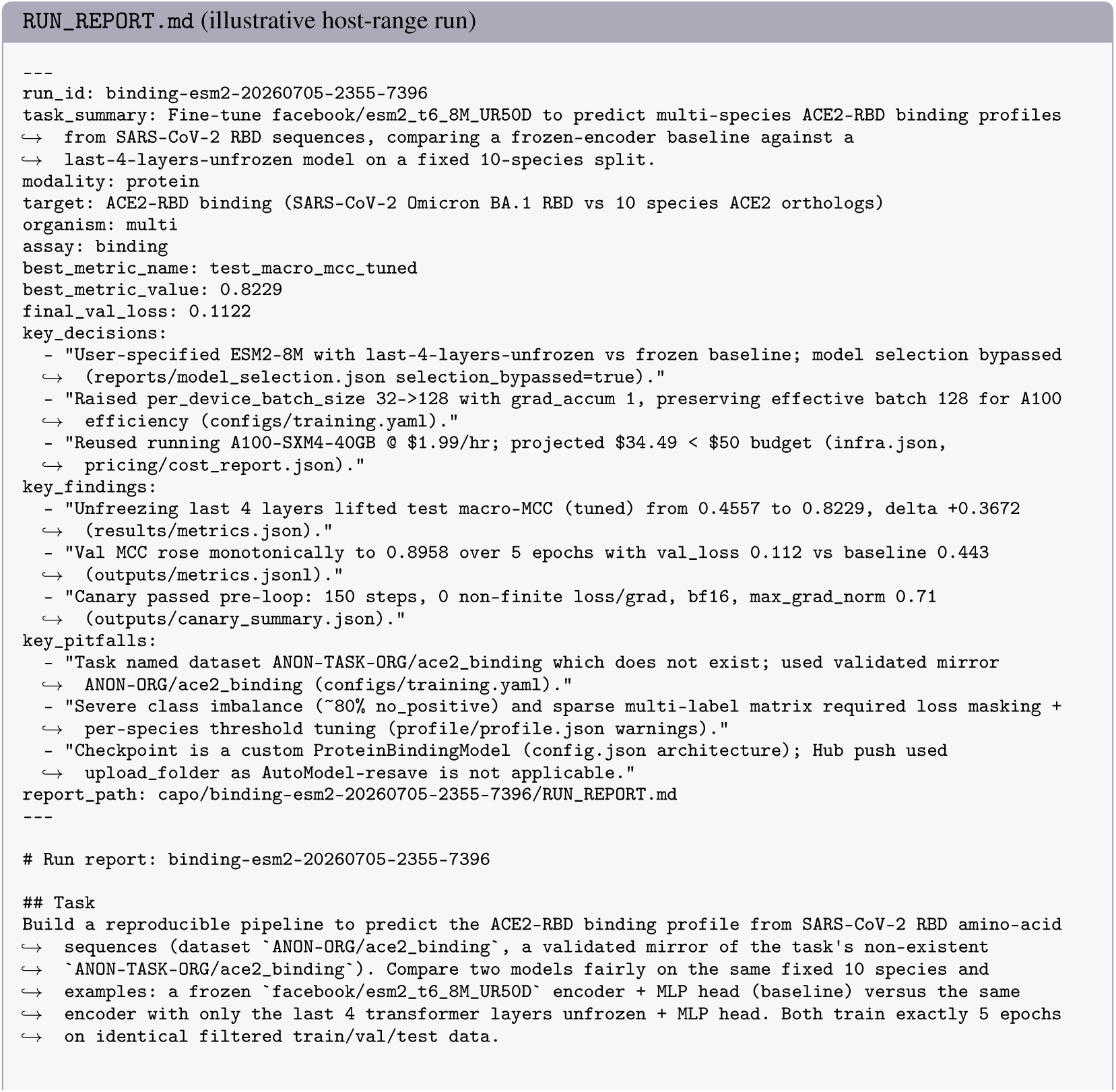
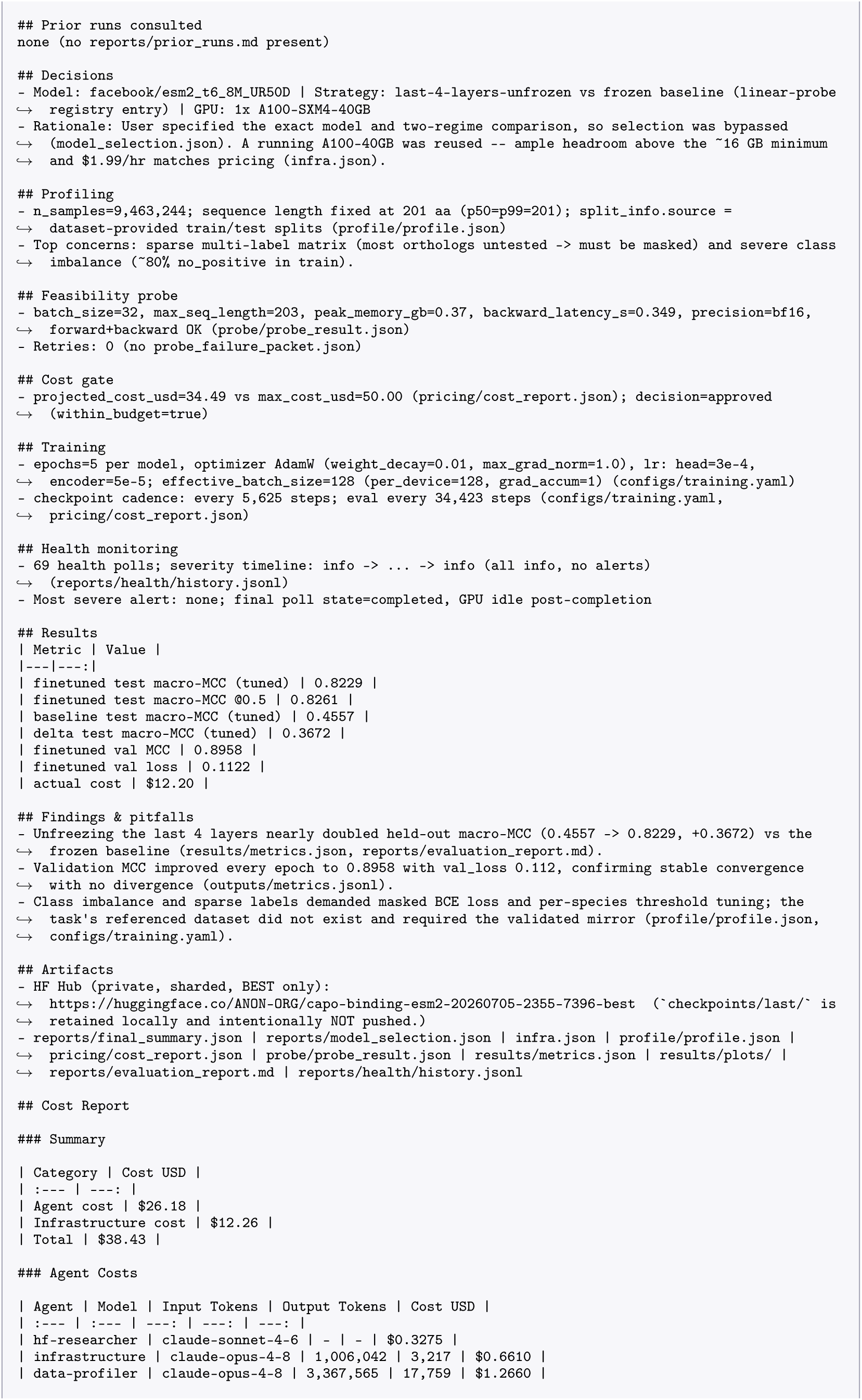
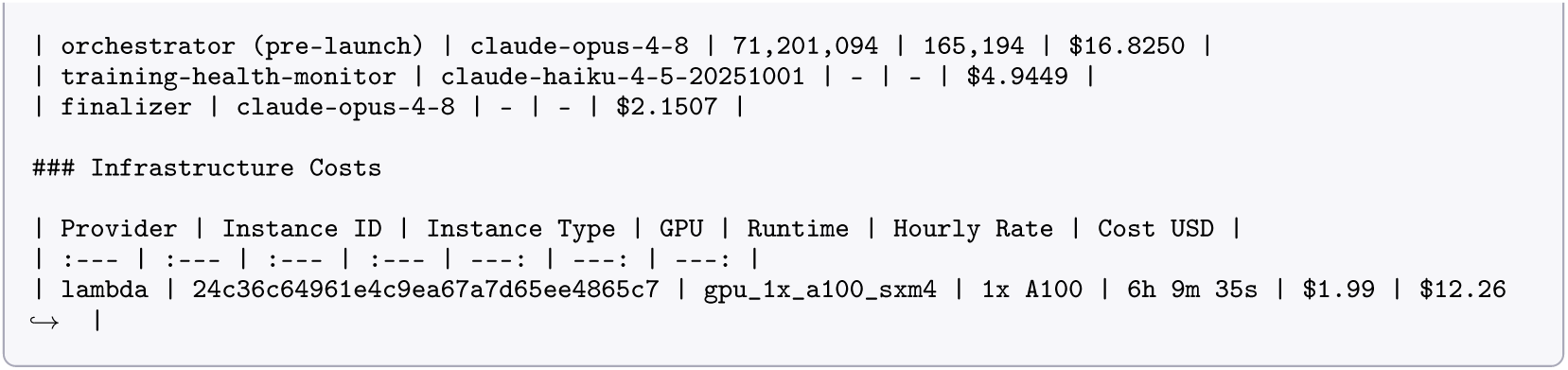

## J User task prompts

Each study began from a single natural-language request, the only task-specific input supplied before the system took over. All three are reproduced verbatim, so that a reader can judge how much the system was told and how much it decided. Dataset and repository identifiers, instance addresses and account names are replaced by placeholders.

### J.1 Host-range binding (ACE2)

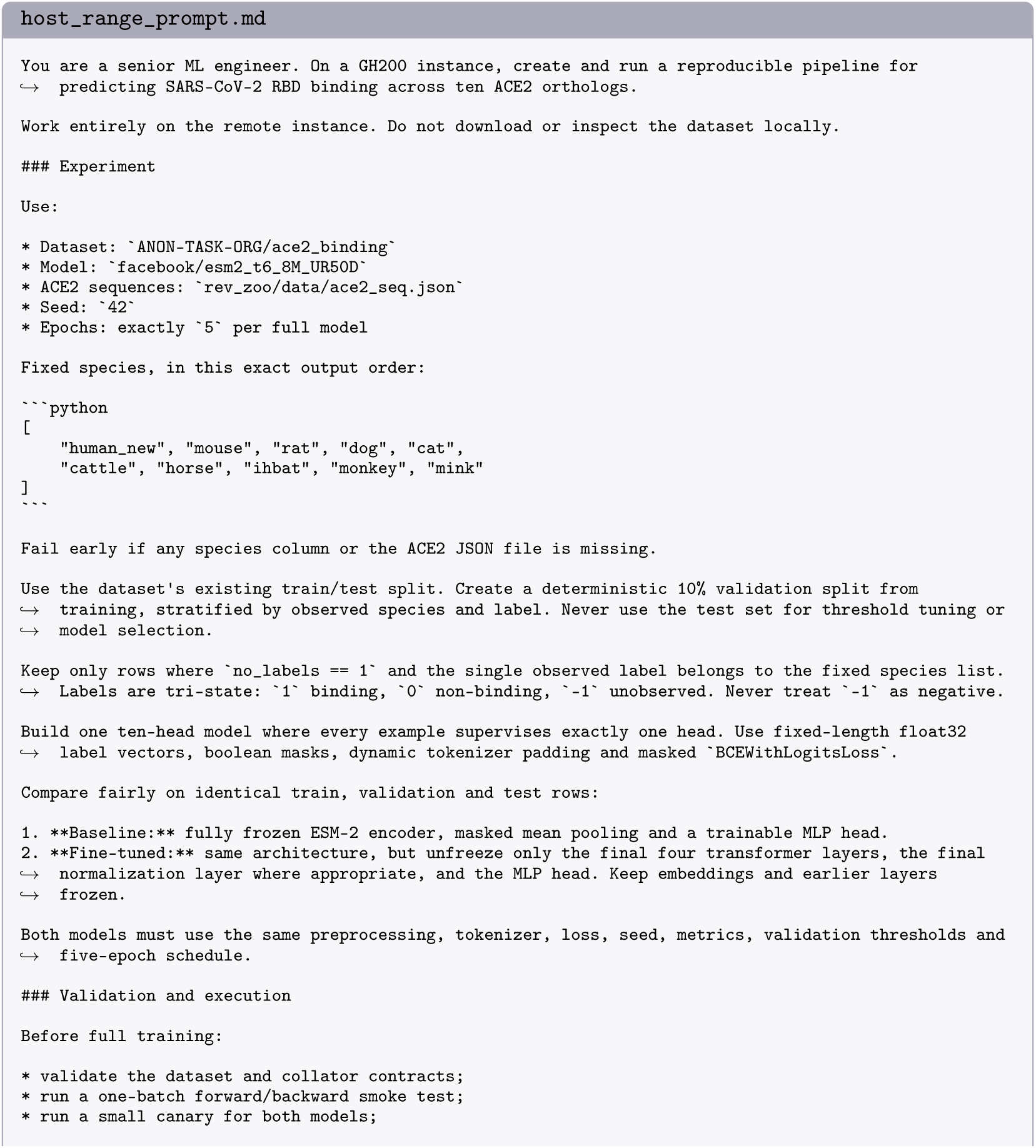
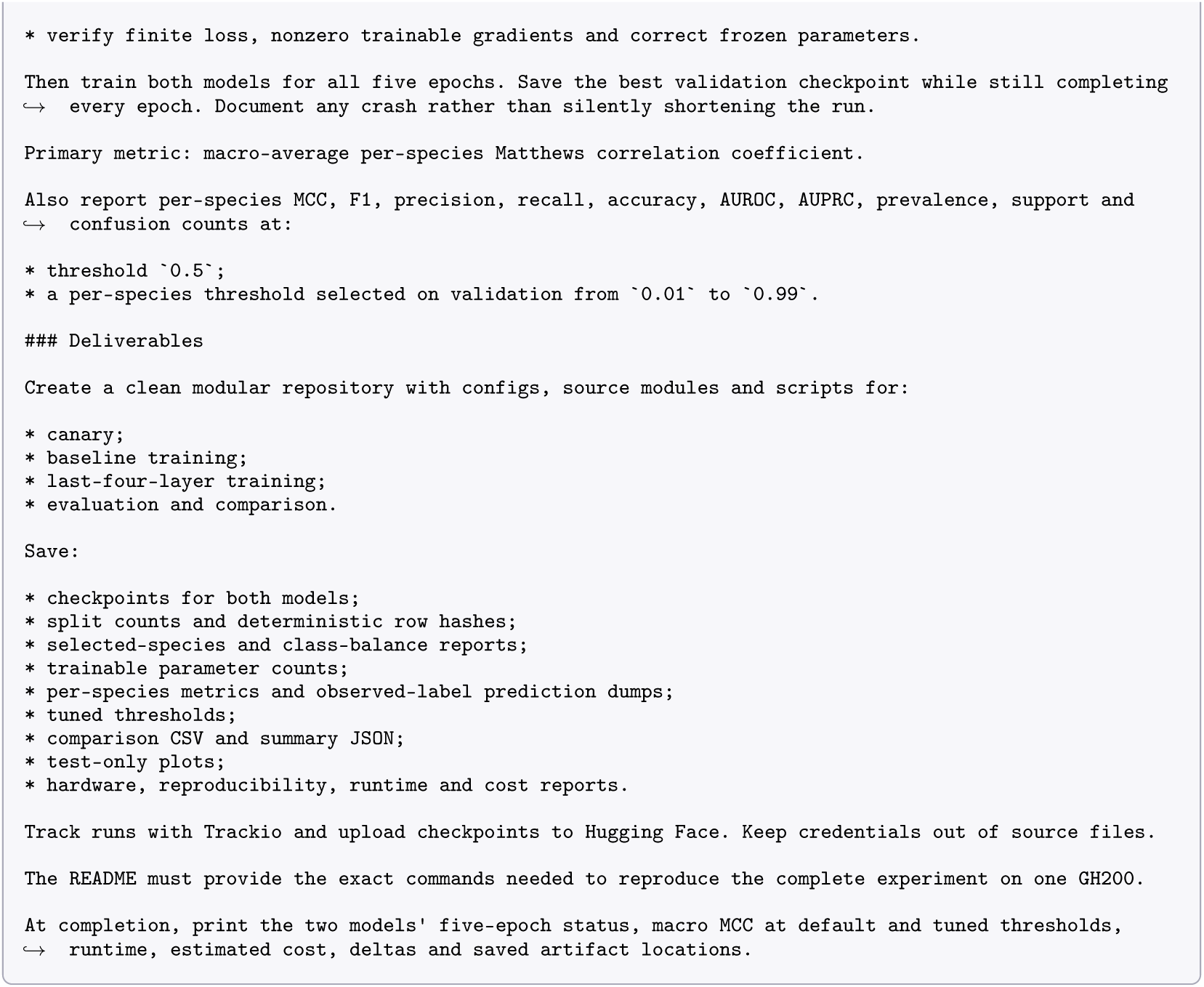

### J.2 Hit-to-lead affinity (Boltz-2)

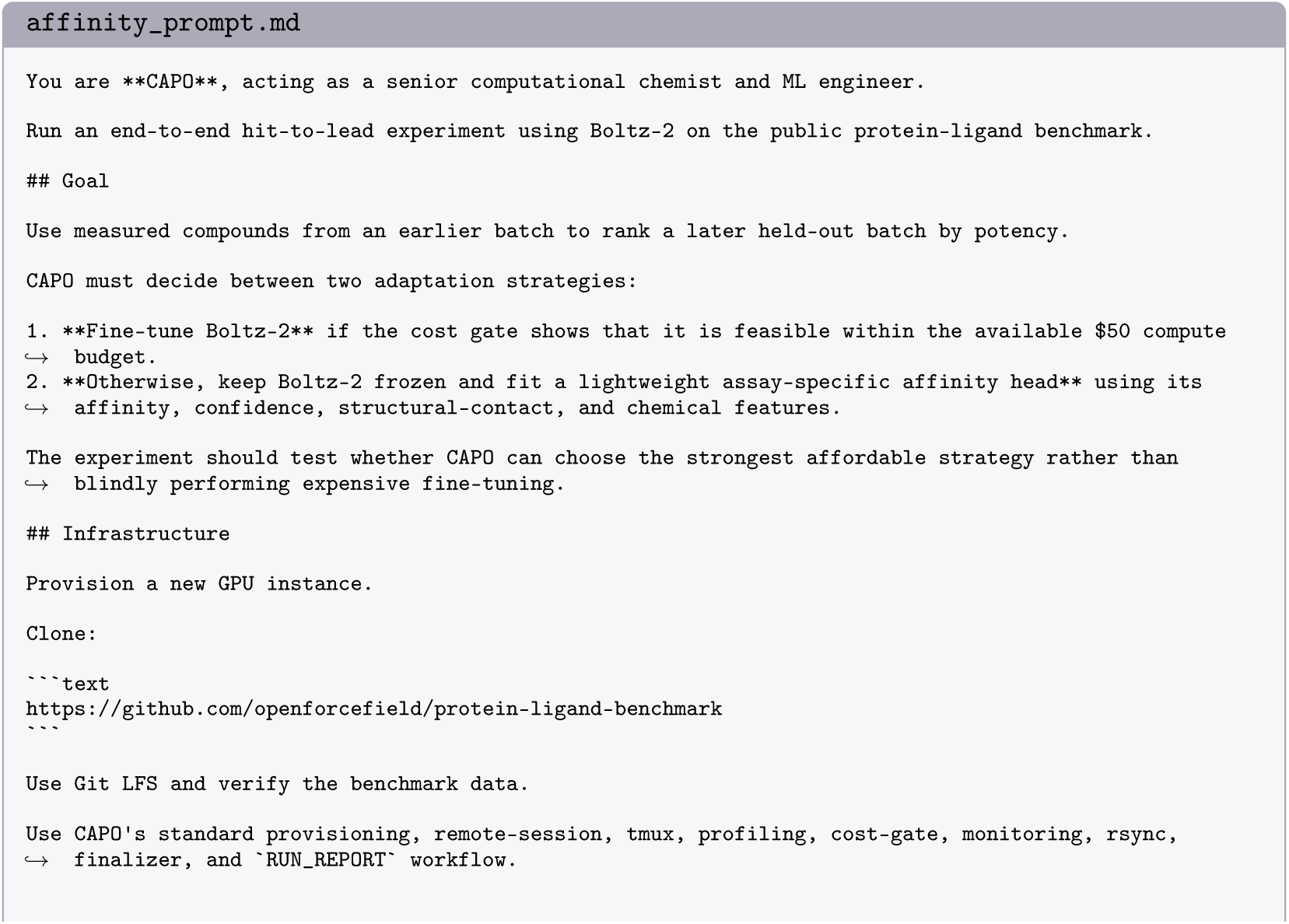
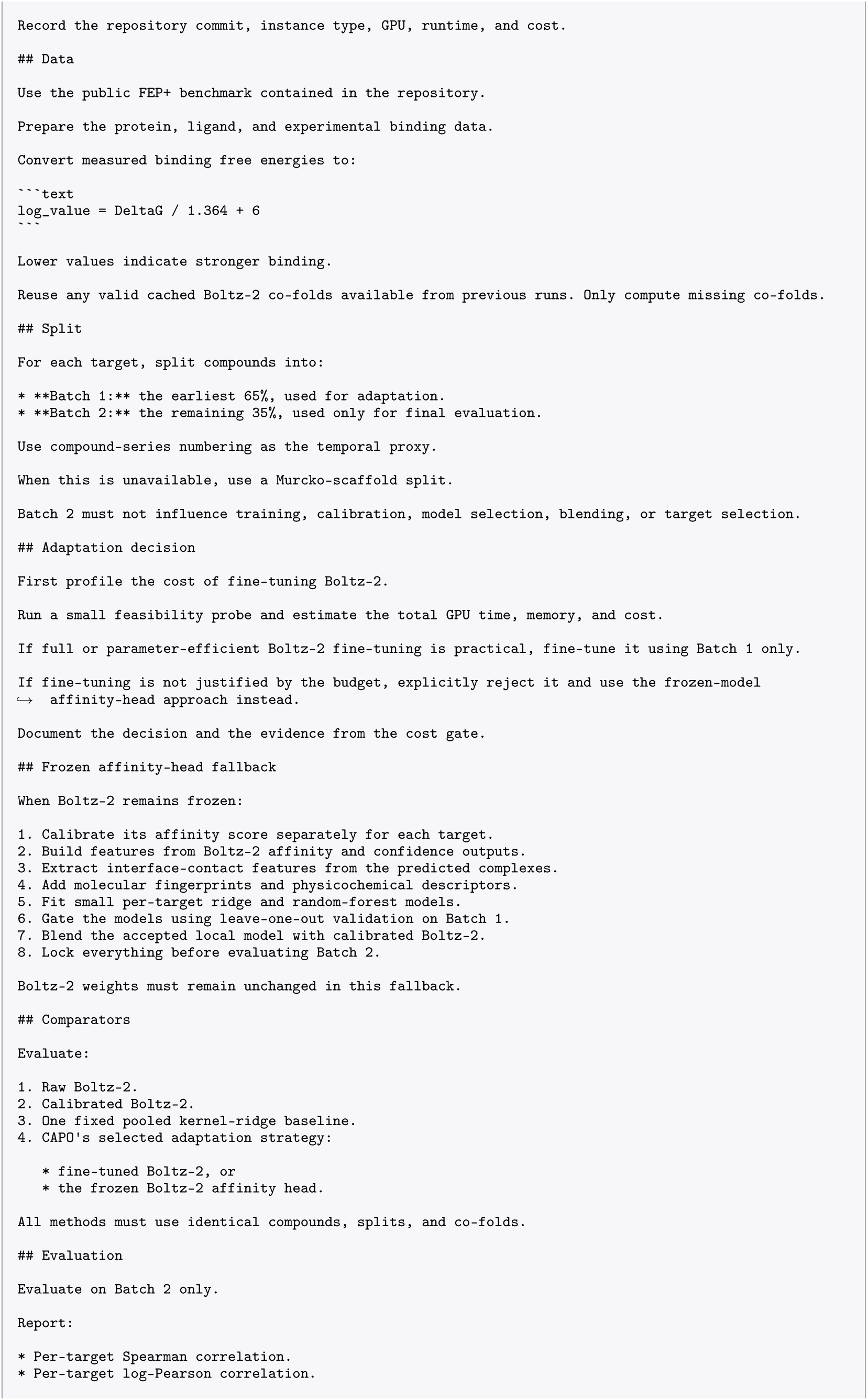
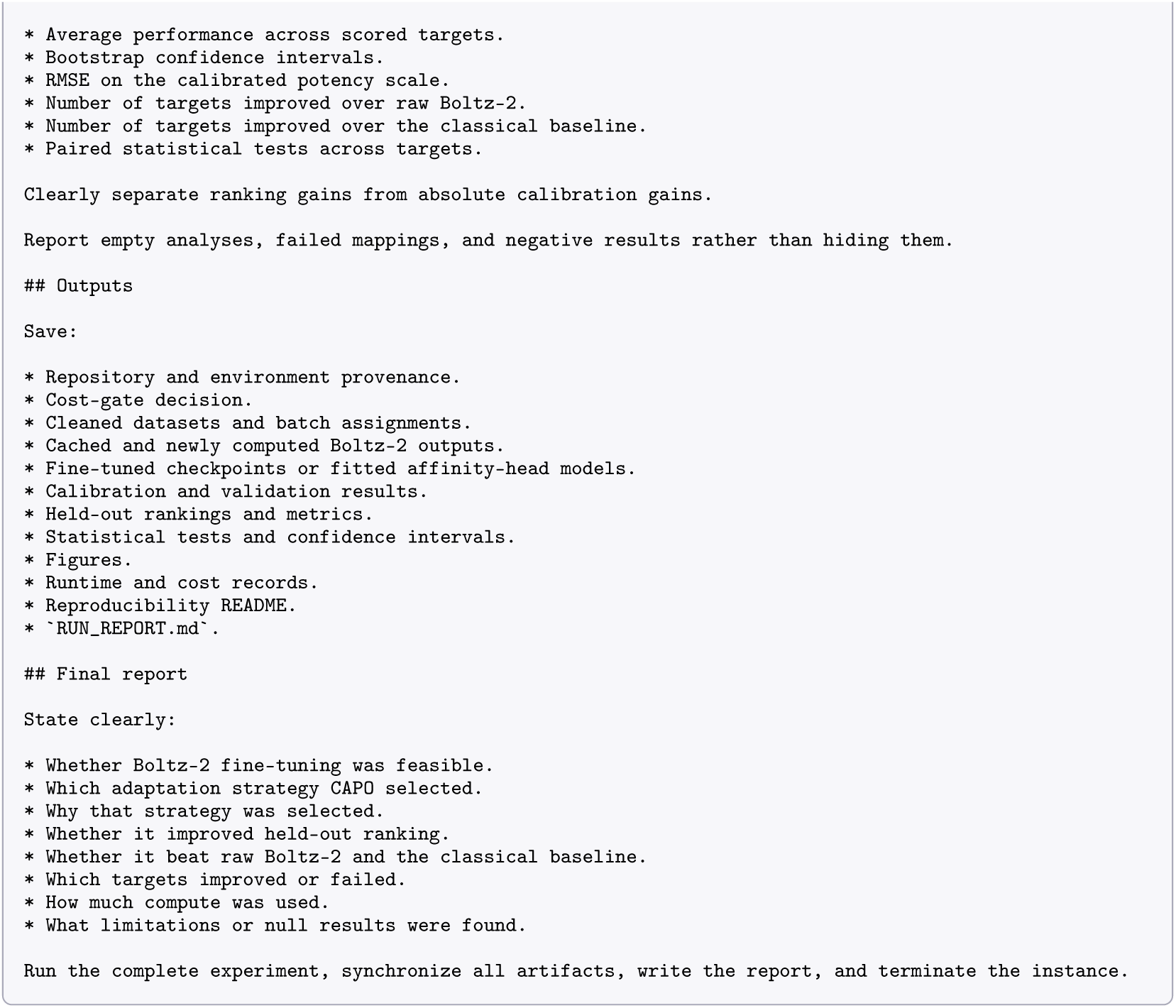

### J.3 Temporal variant effect (ClinVar)

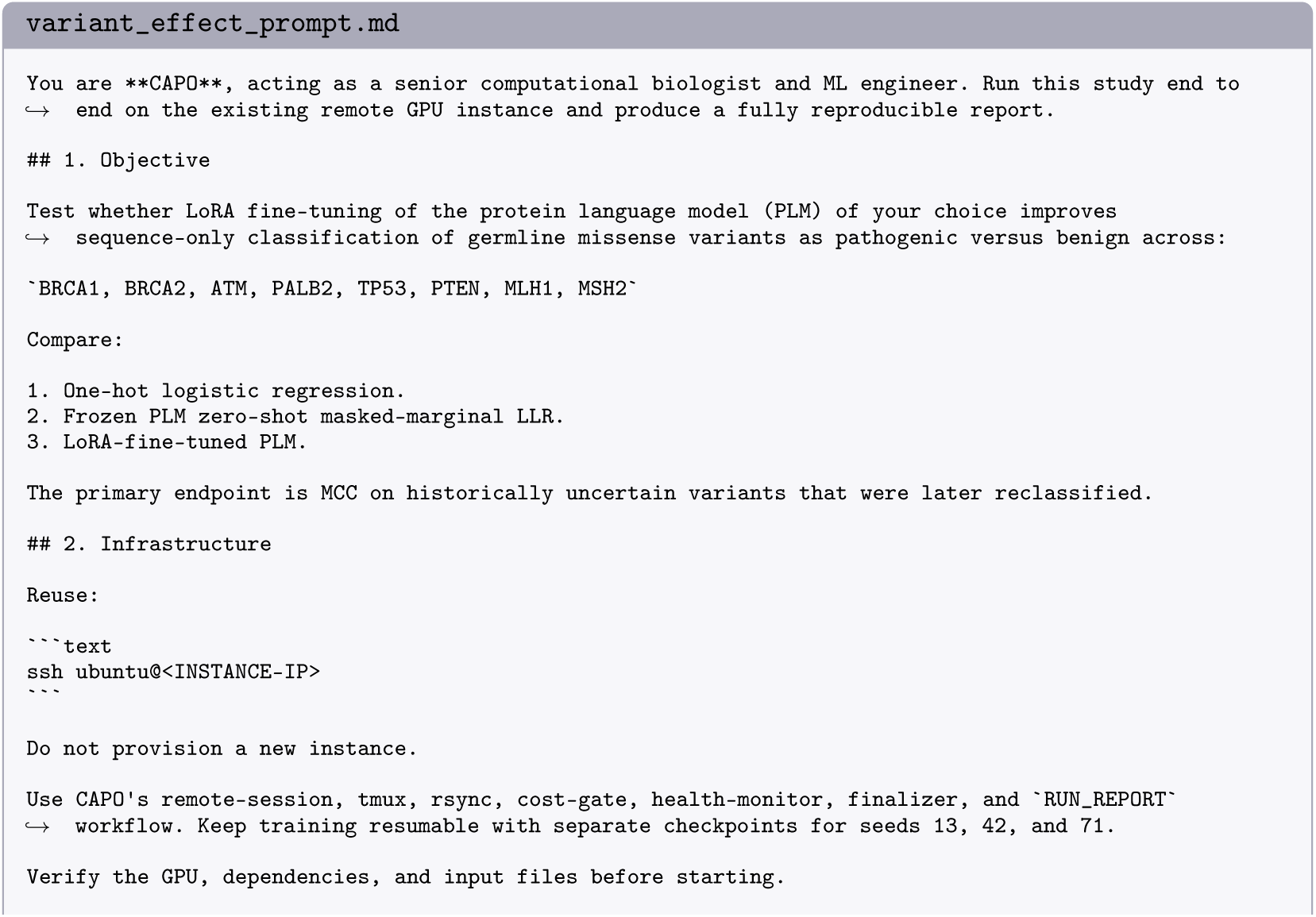
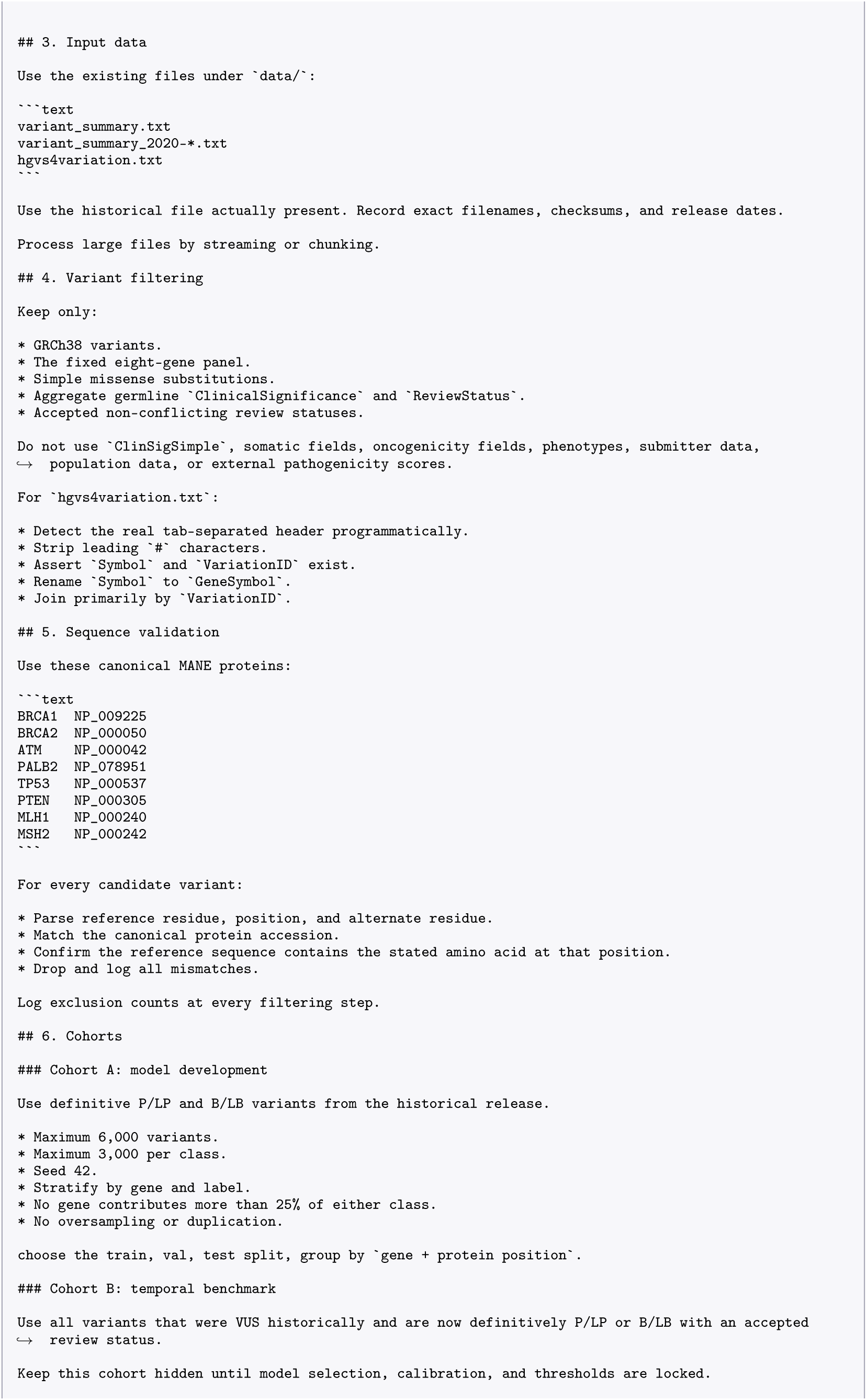
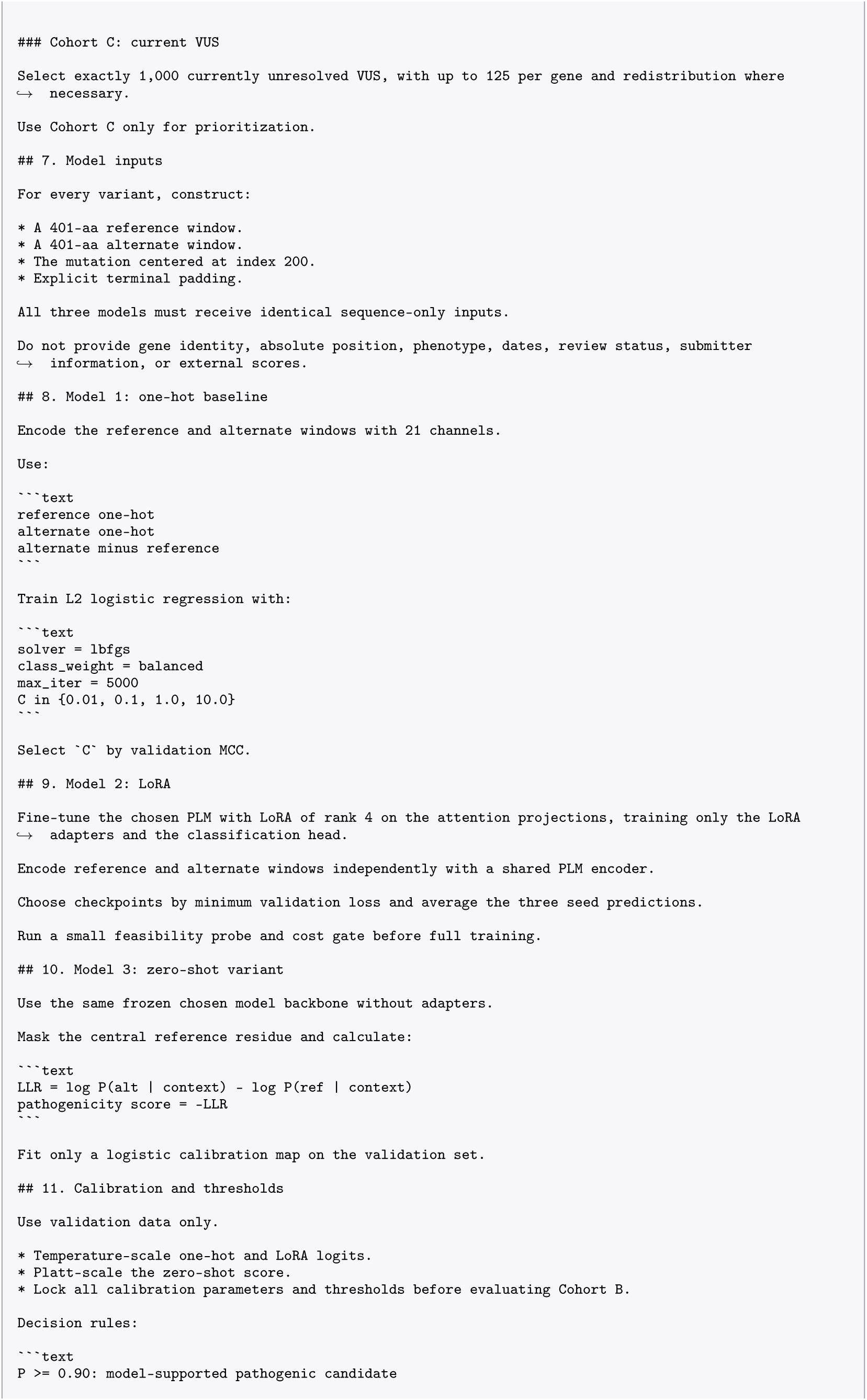
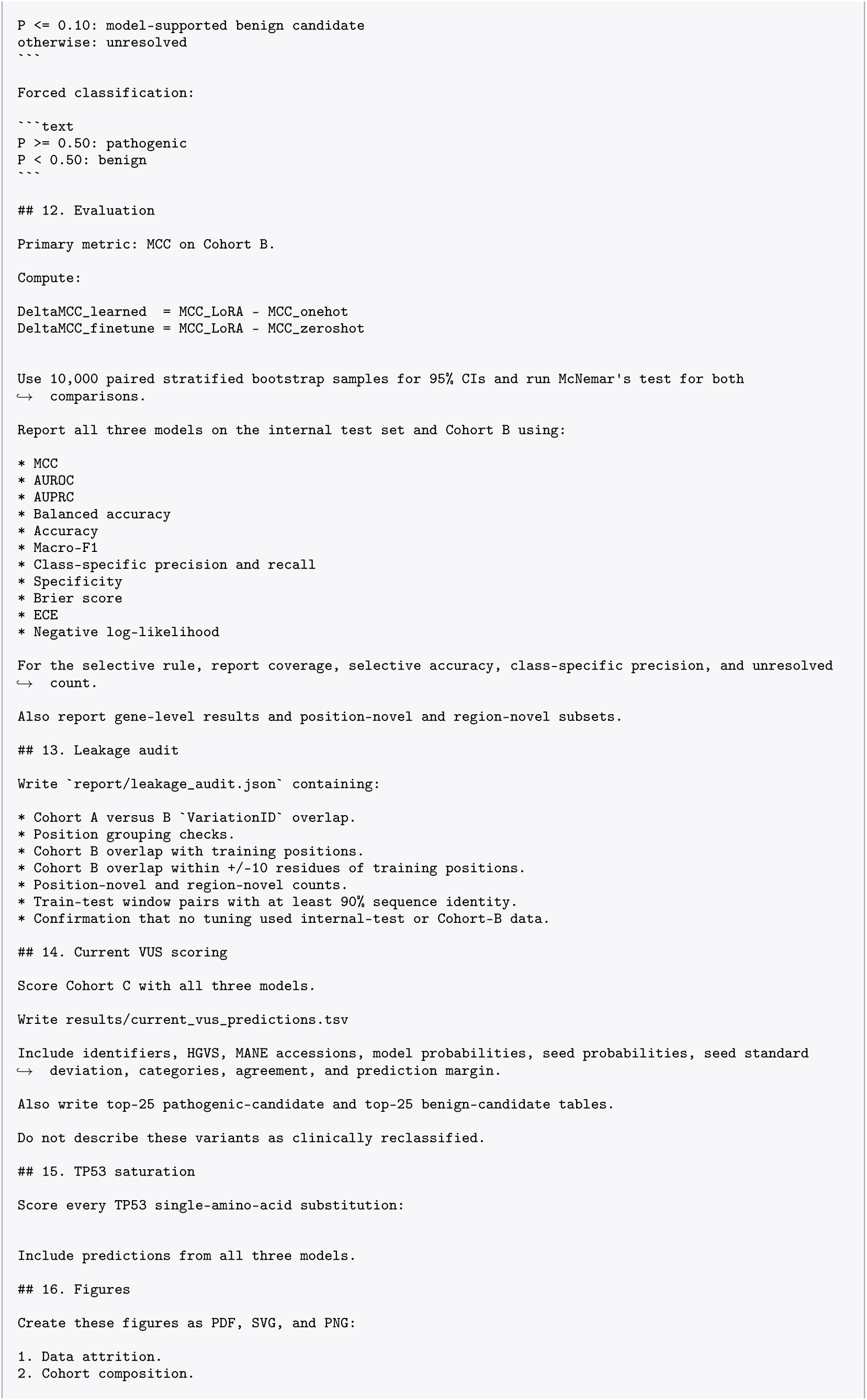
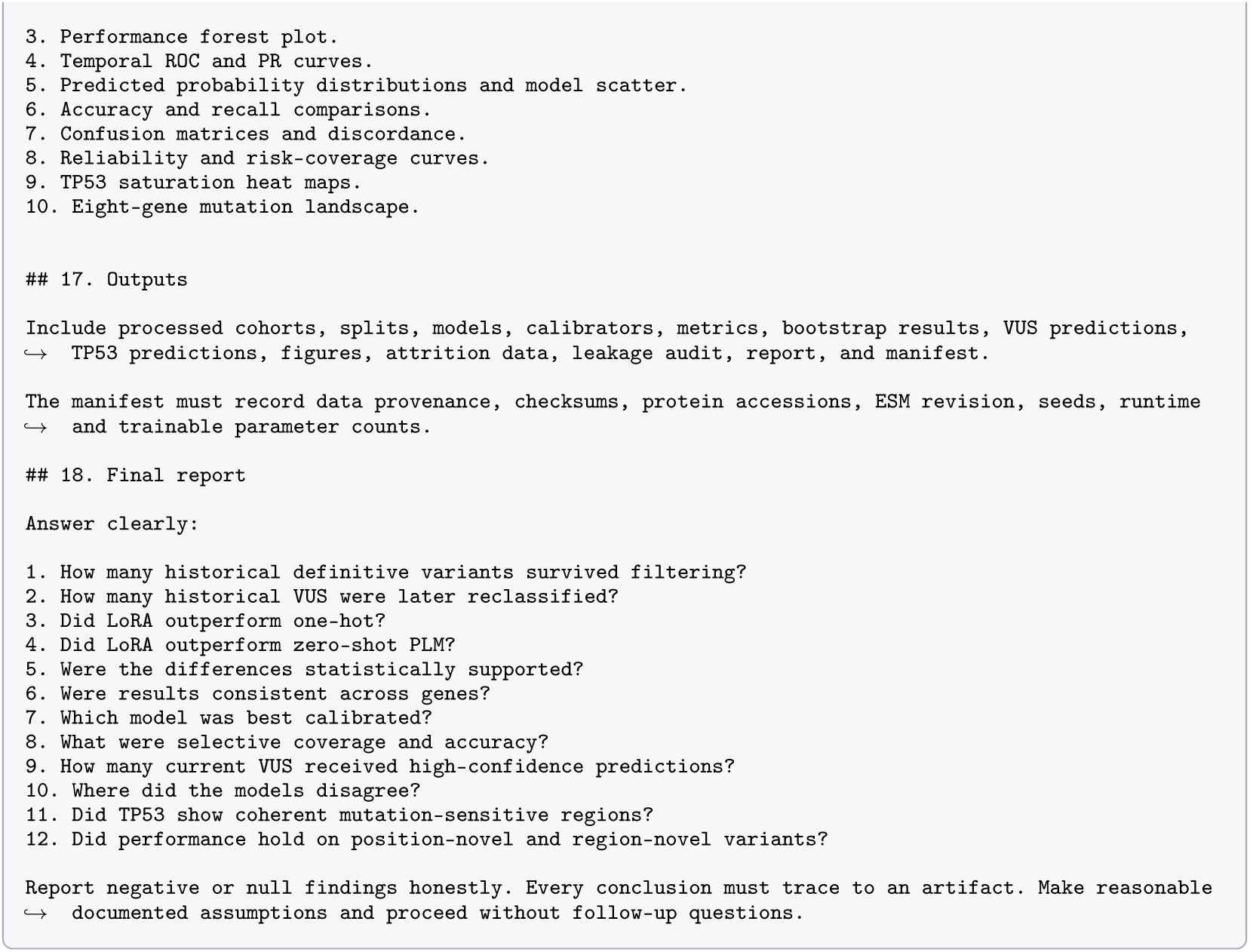

## K Code, data and ethics

The CAPO source, including the verbatim agent system prompts organised by agent, is available at https://github.com/AI-BIIE-Initiative/capo. The hit-to-lead data are the public benchmark at https://github.com/openforcefield/protein-ligand-benchmark; the variant data are the public ClinVar releases at https://ftp.ncbi.nlm.nih.gov/pub/clinvar/tab_delimited/; the host-range assay dataset is available from the authors on request.

### Ethical considerations

This study used public, non-identifiable datasets and privately owned host-range assay data containing no personal or patient information. It involved no direct patient recruitment, clinical intervention, protected health information, patient-level records, medical images or individually linked genomic data. Ethical approval, informed consent and an ethics-committee protocol number therefore did not apply; HIPAA and GDPR requirements specific to identifiable health data were not directly implicated.

The clinical variant task used public ClinVar variant-level annotations. Although some records originate from clinical submissions, CAPO neither attempted identification nor combined records in ways that could enable re-identification. Uncertain variants were used only for research prioritisation and were not presented as clinically reclassified. The model is not intended for diagnosis or clinical decision-making without independent validation and expert review.

Bias remains: ClinVar submissions vary across genes, laboratories, populations and time, and the hereditary-cancer panel spans only eight genes; the binding and affinity data represent specific assays rather than protein biology broadly. CAPO uses temporal and leakage-aware evaluation, class-imbalance reporting, multiple baselines, and preserved uncertainty and negative results. These measures reduce overclaiming but do not eliminate bias or guarantee transfer beyond the evaluated settings.

CAPO supports rather than replaces scientific judgement. Its audit trail preserves evidence, code and run decisions, while feasibility and cost gates reduce unnecessary computation. Human review remains necessary for preprocessing, biological interpretation and any downstream experimental or clinical use.

## Notes

### Competing Interest Statement

The authors have declared no competing interest.

